# Copper-independent lysosomal localization of the Wilson disease protein ATP7B

**DOI:** 10.1101/2022.10.30.514457

**Authors:** Saptarshi Maji, Marinella Pirozzi, Ruturaj, Raviranjan Pandey, Tamal Ghosh, Santanu Das, Arnab Gupta

**Affiliations:** Department of Biological Sciences, Indian Institute of Science Education and Research Kolkata, Mohanpur, West Bengal -741246, India; Institute of Endocrinology and Experimental Oncology, Via P. Castellino, 111, 80131 Napoli, Italy; Children’s Hospital of Philadelphia and University of Pennsylvania, Philadelphia, PA 19104

**Keywords:** ATP7B, Wilson disease, hepatocytes, endolysosomes, Trans-Golgi Network, TGN-proximal lysosomes, TGN-endolysosome contact

## Abstract

In hepatocytes, the Wilson disease protein ATP7B resides on trans-Golgi network and traffics to peripheral lysosomes to export excess intracellular copper through lysosomal exocytosis. We found that in basal copper or even upon copper chelation, a significant amount of ATP7B persists in endolysosomal compartment of hepatocytes but not in non-hepatic cells. These ATP7B-harboring lysosomes lie in close proximity of ∼10nm to the TGN. ATP7B constitutively distributes itself between the sub-domain of the TGN with a lower pH and the TGN-proximal lysosomal compartments. Presence of ATP7B on TGN-Lysosome colocalizing sites upon Golgi disruption suggested possible exchange of ATP7B directly between the TGN and its proximal lysosomes. Manipulating lysosomal positioning significantly alters the localization of ATP7B in the cell. Contrary to previous understanding, we found that upon copper chelation in a copper-replete hepatocyte, ATP7B is not retrieved back to TGN from peripheral lysosomes; rather, ATP7B recycles to these TGN-proximal lysosomes to initiate the next cycle of copper transport. We report a hitherto unknown copper-independent lysosomal localization of ATP7B and the importance of TGN-proximal lysosomes but not TGN as the terminal acceptor organelle of ATP7B in its retrograde pathway.

**Synopsis:** Excess copper is toxic to the cell. In hepatocytes, the Wilson disease protein ATP7B has been reported to recycle between the trans-Golgi network and lysosomes to export copper in elevated copper conditions. We, for the first time, show that a large fraction of ATP7B constitutively resides on the lysosome and rather, the position of the lysosome shifts from TGN-proximal to the peripheral region of the cell in high copper to facilitate ATP7B-mediated lysosomal exocytosis of copper.

## Introduction

Wilson disease (WD) is caused due to mutations in the copper-transporting ATPase leading to copper accumulation in the liver ^1–5^. Lysosomal exocytosis serves as a key process in order to maintain optimal intracellular levels of the physiologically crucial micronutrient, copper ^6–9^. Lysosome is the primary organelle that serves as the regulatory hub for recycling and homeostasis in a cell. Being a dynamic organelle, it interacts with other organelles and endosomal system to maintain a multitude of crucial homeostatic functions ^10–12^. A variety of cellular signalling can affect lysosomal size, number, and position rapidly. The distribution of lysosomes in terms of its size, position, acidity, and number are important indicators of cellular health. These factors are connected and account for the proper functioning of the lysosome. Lysosomal exocytosis is one of the many important processes that lysosome undergoes to serve several important purposes like cellular clearance, membrane repair, and secretory functions. Toxic cellular disposals including pathogens, heavy metals, etc., that are deposited to the lysosome are cleared by this essential process, and failure to do this can lead to several pathological conditions ^13^.

Lysosomal positioning is another important factor depicting lysosome health related to lysosomal exocytosis ^14^. Depending on the distance from the nucleus, lysosomes can be divided into two types. Lysosomes near to nucleus are known as perinuclear lysosomes. Lysosomes away from the nucleus are known as peripheral lysosomes ^15^. These two types of lysosomes are different from each other in their luminal content and are tightly regulated. mTOR is one of the important regulators of these lysosomes. mTOR activation leads to anterograde trafficking of lysosome toward the plasma membrane, whereas mTOR downregulation leads to retrograde trafficking of lysosome toward nucleus ^16,17^.

Copper is an important micronutrient for all eukaryotes, acting as a co-factor for an array of cupro-enzymes. But copper, at an elevated level, is detrimental to the cell. Hence the concentration of intracellular copper is tightly regulated ^18^. Copper is up-taken in the cell via high-affinity copper transporter CTR1, where environmental Cu^2+^ is reduced to bio-available and utilizable Cu^+^^19,20^. Subsequently, copper is sequestered by cellular copper chaperones that deliver copper to different parts of the cell. Atox1 delivers copper to TGN-localized ATP7A and ATP7B, the two copper ATPases ^21^. At the physiological copper level, ATP7A and ATP7B pump copper into the TGN lumen that serves as a site for the maturation of cuproproteins. At elevated cellular copper condition, the Cu-ATPases vesicularizes from TGN and traffics to endolysosomes to export out excess copper ^22^

Several factors govern the trafficking of ATP7B. The cytoplasmic amino-terminal sequence was found to be important for copper-responsive trafficking of ATP7B ^23,24^. WD-causing mutations on conserved cytoplasmic domain of ATP7B can lead to either loss of copper-responsive trafficking from the TGN or constitutive vesicularization of the protein, even in copper-limiting conditions. Engineered mutation on the tri-leucine motif (^1454^LLL^1456^) causes constitutive vesicularization of the protein ^25^. Similarly, S340/341A mimics copper saturated amino terminal of ATP7B leading to copper-independent vesicularization ^23^. On the other hand, high copper fails to trigger the trafficking of ATP7B, harbouring the WD-causing mutation S653Y ^26^. Recent studies have shown that ATP7B can even traffic without the influence of copper in presence of several anti-cancer drugs ^27^.

The crosstalk between lysosome and ATP7B is important for maintaining proper copper homeostasis inside the cell. With increasing cellular copper levels, ATP7B traffics toward endo-lysosomal compartment to pump the excess copper into the lysosomal lumen. Increase in cellular copper level downregulates mTOR activity, which in turn activates TFEB mediated lysosomal biogenesis ^28^. Elevated copper also triggers lysosomal exocytosis, and thus excess copper is excreted out ^6,7,29^.

In this study, we have demonstrated that a significant fraction of the total ATP7B pool is present on the endo-lysosomal compartment irrespective of cellular copper concentration and this localization of ATP7B is exclusively hepatocyte-specific. We further observed a pool of lysosomes apposing close to the TGN that harbours this fraction of ATP7B in basal or copper chelated condition. We hypothesise presence of a novel organelle contact site between TGN and lysosome/endolysosomal compartments. We also found that lysosomal positioning influences localisation of ATP7B at these TGN-proximal lysosomes. Finally, we have demonstrated that the recycling of ATP7B by the retrograde pathway that is triggered by copper chelation in copper replete cells, terminates at TGN-proximal lysosomes in contradiction to TGN as the terminal location as previously reported.

## Results

### ATP7B localizes on endolysosomes independent of intracellular copper concentration

Liver is the major site for copper detoxification in mammals. The Wilson disease protein, ATP7B is sole regulator of physiological copper levels in the mammalian liver. ATP7B is primarily expressed in hepatocytes ^30^, though its expression in non-hepatic tissues is also reported ^7,31,32^. It is well documented that under basal intracellular conditions ATP7B localizes at the trans-Golgi network (TGN) and helps in transporting copper to the TGN lumen where it is utilized by various copper dependent proteins ^33^. In hepatocytes, at elevated intracellular copper ATP7B traffics to the endolysosomal compartment for exporting copper from cytosol which is subsequently excreted out of the cell via copper-induced increased lysosomal exocytosis ^1,3,34^. Upon removal of copper by chelators, ATP7B has been reported to recycle back to the TGN ^35^.

We determined localization of ATP7B in HepG2 cells in basal and elevated copper conditions. Interestingly, to our surprise at basal copper (cells not treated with extracellular copper), a significant amount of ATP7B constitutively localized at the endolysosomal compartment as determined its colocalization with the lysosomal marker LAMP2 (Fig.1A). We found that 56.4±21% (mean±SD) of total ATP7B localizes at the endolysosomal compartments in basal conditions. As expected, upon copper treatment (20, 100 and 250µM), the fraction of ATP7B at the TGN decreases coupled with its increased colocalization with LAMP2 (Fig.S1A, quantitated in Fig.1B). When copper was chelated from the cell using cell-impermeable copper chelator BCS (Bathocuproinedisulfonic acid); 100 µM for 2h or cell permeable chelator TTM (Tetrathiomolybdate); 25µM for 2h, the endolysosomal localization of ATP7B diminish but do not entirely disappear (Fig.1A, S1B; quantitated in Fig.1B). We found a similar trend in colocalization of ATP7B with another endolysosomal marker, LAMP1 (Fig.S1C, S1D).

**Figure 1:**
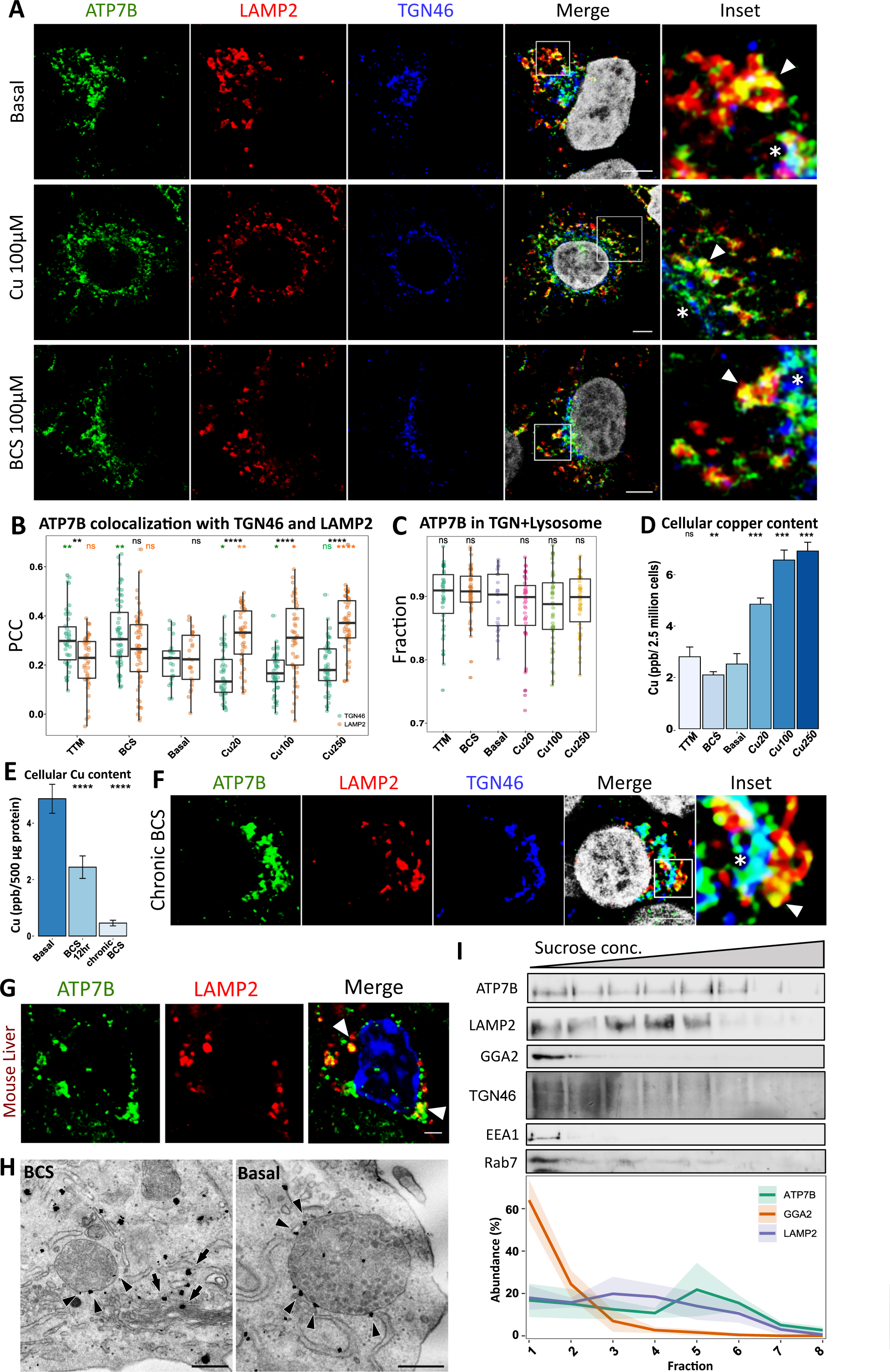
ATP7B distribution at the TGN and endolysosomes in different copper treatments. **(A)** Immunofluorescence image of ATP7B (green) in HepG2 cell-co-stained with endolysosomal marker LAMP2 (red) and TGN marker TGN46 (blue) shows presence of ATP7B on both compartments, irrespective of cellular copper condition i.e., basal, high copper (100µM Cu) as well as copper chelated conditions (100µM BCS). The marked area on ‘merge’ panel is enlarged in ‘inset’. The arrowhead shows colocalization between ATP7B-LAMP2, the asterisk shows colocalization between ATP7B-TGN46. **(B)** Pearson’s Colocalization Coefficient (PCC) between ATP7B-TGN46 and ATP7B-LAMP2 in different copper conditions are demonstrated by box plot with jitter points. The black * depicts the comparison of colocalization between ATP7B-TGN46 and ATP7B-LAMP2. The green * shows the comparison of ATP7B-TGN46 colocalization under different copper conditions w.r.t. basal condition. The orange * shows the comparison of ATP7B-LAMP2 colocalization under different copper conditions w.r.t. basal condition. **(C)** Cumulative fraction of ATP7B on TGN46 and LAMP2 positive compartment are demonstrated by box plot with jitter points. The statistical comparison is done w.r.t. Basal. Sample size (N) for (B) and (C) are TTM: 42, BCS: 57, Basal: 24, Cu20: 44, Cu100: 49, Cu250: 50. **(D)** comparison of copper accumulation in HepG2 cells (n = 9) under different treatment conditions shows elevated copper accumulation in case of increased copper treatment, copper concentrations were measured in parts per billion (ppb). Data is demonstrated as bar-plot (mean ± SD). Statistical comparison is done w.r.t basal. **(E)** Comparison of copper accumulation in HepG2 cells (n = 9) under prolong copper chelated conditions (12h as well as 72h chronic chelation using BCS) shows decreased copper accumulation. Copper concentrations were measured in parts per billion (ppb). Data is demonstrated as bar-plot (mean ± SD). **(F)** Immunofluorescence image of ATP7B (green) in HepG2 cell line co-stained with endolysosomal marker LAMP2 (red) and TGN marker TGN46 (blue) shows presence of ATP7B on both the compartments even under chronic copper chelated condition. The marked area on ‘merge’ panel is enlarged in ‘inset’. The arrowhead shows colocalization between ATP7B-LAMP2, the asterisk shows colocalization between ATP7B-TGN46. **(G)** Immunofluorescence image of ATP7B (green) co-stained with LAMP2 (red) in mouse tissue shows presence of ATP7B on LAMP2 positive compartment (marked by arrowhead). **(H)** Immuno-EM of endogenous ATP7B in HepG2 cell shows presence of ATP7B at the Golgi membrane (marked by arrow) as well as late-endosome/lysosome compartment (marked by arrowhead) both in BCS treated (left panel) and basal (right panel) conditions. **(I)** Immunoblot of ATP7B, lysosomal marker LAMP2, TGN marker GGA2 and TGN46, early endosome marker EEA1 and late endosome marker Rab7 of the top 8 fractions of sucrose gradient shows presence of ATP7B on TGN and lysosome enriched fraction. The line plot shows abundance of ATP7B (green), GGA2 (orange) and LAMP2 (violet) in 8 fractions of sucrose gradient. The ribbon plot signifies the standard deviation of 4 independent experiments. [scale bar for EM: 100 nm, for immunofluorescence: 5µm]

Interestingly, the combined fraction of ATP7B present in TGN and endolysosomal compartment remains unchanged irrespective of copper treatment supporting the idea of ATP7B distributes exclusively between these two compartments (Fig.1C). Intracellular copper concentration with treatment or chelation in the media was confirmed by ICP-OES measurement of cellular copper concentration (Fig.1D). Observing that levels of intracellular copper did not significantly decrease in cells treated with 100µM BCS or 25µM TTM for 2h (Fig.1D), leave open a possibility that residual copper in those sets were enough to trigger lysosomal localization of ATP7B. To further deplete copper, we maintained the cell for 72h (3 days) in 10µM BCS (chronic treatment) followed by 2h of 100µM of BCS. In a second set, we treated the cells with 100µM BCS for 12h. Upon measuring copper, we recorded a drop of ∼90% and 50% copper respectively as compared to cells maintained in basal media (Fig. 1E). Interestingly, in agreement with our previous observations, a fraction of ATP7B persisted on the lysosomes in close apposition with the TGN (Fig.1F, Fig.S1E), the colocalization of which did not seem to vary between cells in basal media with cells under severe copper chelation (Fig.S1F). Total abundance of the ATP7B protein did not change significantly with varying copper levels in the treatment media, though a slight increase was observed for cell treated with 100µM and 250µM copper (Fig. S1G). In liver sections of mice where we also observed that in basal conditions, a fraction of ATP7B colocalized on lysosomes marked with LAMP2 (Fig.1G).

Using immune-EM, we further achieved a higher resolution and observed that indeed ATP7B decorated the lysosomal membranes, besides being resident on the TGN in basal as well as copper chelated conditions (Fig.1H).

Further to biochemically substantiate our finding on distribution of ATP7B in lysosome vs TGN, we employed sub-cellular fractionation to isolate 8 fractions of cellular lysate and identified different Golgi, endosomal and endolysosome rich fractions. Presence of ATP7B in gradient fractions positive for GGA2 and LAMP2 confirms its localization at the TGN as well as lysosomes respectively in basal copper conditions. (Fig.1I). However, the gradual disappearance of GGA2 from the TGN fractions beyond fraction 2 might be attributed to the fact that it being a peripheral membrane protein, could potentially dissociate from the membranes during the sucrose gradient process. To circumvent this potential problem, we utilized TGN46 as a TGN marker to confirm the presence of ATP7B in the Golgi. Remarkably, the peak corresponding to fraction 5 that shows highest ATP7B signal moderately overlaps with the endolysosomal marker LAMP2 that interestingly shows maximum abundance at fraction 3-4. This is probably because LAMP2 marks various phases and types of lysosomal-related organelles and ATP7B is associated with a specific type of it, that is still to be characterized ^36^. However, we failed to notice any appreciable presence of early endosomal marker, EEA1 and late endosomal marker, Rab7 in the gradient fractions of ATP7B (Fig.1I). Lack of colocalization between ATP7B with EEA1 or Rab7 in immunofluorescence assay further validated our findings (Fig.S1H and S1I).

Since, hepatocytes deploy lysosomal exocytosis to export excess copper out of the cell, we found our observation of ATP7B localizing in lysosomal compartments even in basal and copper chelated conditions thought provoking that demanded further investigation.

### Endolysosomal localization of ATP7B in basal copper condition is hepatocyte-specific

We further investigated whether this property of ATP7B localizing at endolysosomal compartments in low and limiting copper is a ‘cell-type’ specific phenomenon or is observed globally. In Huh7, another hepatic cell line derived from hepatoma tissue, we found a similar localization pattern of ATP7B (Fig 2A, 2B). Levels of copper did not significantly influence the abundance of total ATP7B in these cells as observed earlier in HepG2 (Fig. 2C). To further examine our findings in non-hepatic cells, we utilized HeLa and kidney cell line HEK293T. Presence of ATP7B in other cell lines were confirmed by immunoblotting and compared with hepatic cell lines, HepG2 and Huh7 (Fig 2D). Interestingly, both in HeLa and HEK293T, endogenous ATP7B were present exclusively in TGN in basal copper condition with no endolysosomal localisation. However, as observed in hepatocytes, in elevated copper condition ATP7B traffics from TGN to endolysosomal compartment in HeLa that was determined by diminished colocalization of ATP7B with TGN46 and increasing colocalization with LAMP2 (Fig. 2E, 2F). Interestingly, in HEK293T cells, as previously reported, minimal TGN exit of ATP7B was observed in high copper condition as determined by continued colocalization between ATP7B and TGN46 (Fig S2A). To examine if the difference in ATP7B localization may be due to underlying sequence differences of the cDNA between hepatic and non-hepatic cells, we ectopically expressed mEGFP-ATP7B (cloned from liver cDNA) in HeLa. The expressed protein also did not exhibit any endolysosomal localization at basal or copper-limiting condition. At elevated copper concentration, mEGFP-ATP7B was found to be exiting TGN and localizing at the endolysosomal compartment which confirms copper responsive trafficking of ATP7B in non-hepatic cell lines (Fig S2B).

**Figure 2:**
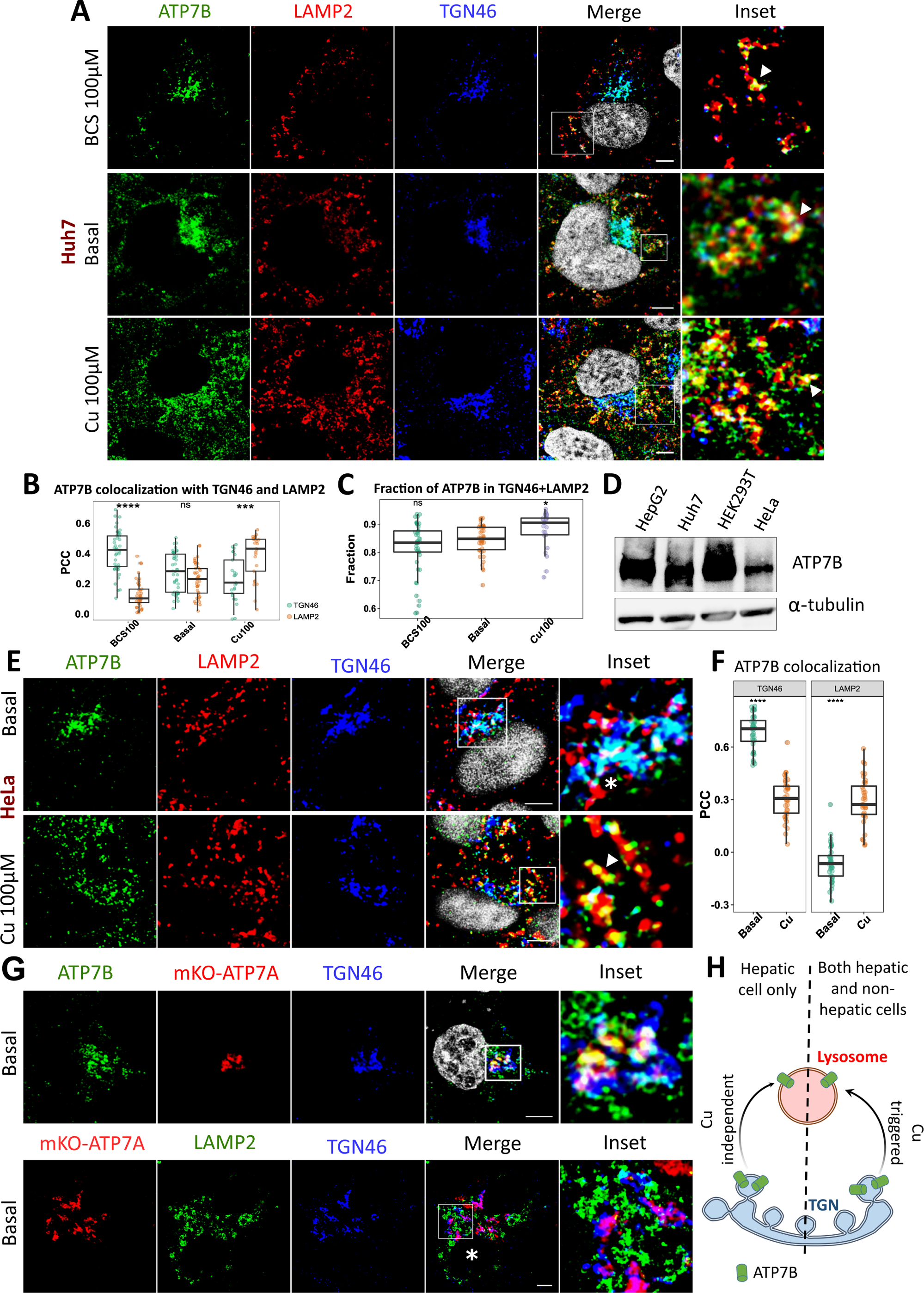
Distribution of ATP7B and ATP7A in hepatic and non-hepatic cell lines. **(A)** Immunofluorescence image of ATP7B (green) in Huh7 cell line co-stained with LAMP2 (red) and TGN46 (blue) shows presence of ATP7B on both the compartments, irrespective of cellular copper conditions. The marked area on ‘merge’ panel is enlarged in ‘inset’. The arrowhead shows colocalization between ATP7B-LAMP2. **(B)** The PCC between ATP7B-TGN46 and ATP7B-LAMP2 in different copper condition in Huh7 cells are demonstrated by box plot with jitter points. **(C)** Total fraction of ATP7B present in TGN46 and LAMP2 compartment are demonstrated by box plot with jitter points. The statistical comparison is done w.r.t. basal. Sample size (N) for (B) and (C) are BCS: 45, Basal: 42, Cu100: 25. **(D)** Immunoblot of cell lysate shows presence of endogenous ATP7B (∼165kDa) in different cell lines. α-tubulin (∼55kDa) has been used as housekeeping protein. **(E)** Immunofluorescence image of ATP7B (green) in HeLa cell line co-stained with LAMP2 (red) and TGN46 (blue) shows different localization pattern of ATP7B that hepatocytes. The marked area on ‘merge’ panel is enlarged in ‘inset’. The arrowhead shows colocalization between ATP7B-LAMP2, the asterisk shows colocalization between ATP7B and TGN46. **(F)** The PCC between ATP7B-TGN46 and ATP7B-LAMP2 in different copper condition are demonstrated by box plot with jitter points. Sample size (N) for (E) are Basal: 36 Cu: 36. **(G)** Ectopically expressed mKO-ATP7A (red) in HepG2 co-stained with ATP7B (green) and TGN46 (blue) (*top panel*) and LAMP2 (green) and TGN46 (blue) (*bottom panel*) shows distinct localization pattern of ATP7A than ATP7B. The marked area on ‘merge’ panel is enlarged in ‘inset’. Asterisk marks the nucleus of the cells in the bottom panel. **(H)** Model illustrates hepatocyte-specific copper-independent, and copper-dependent lysosomal localization of ATP7B. [Scale bar: 5µm.]

We tested if the localization pattern of ATP7B in hepatocytes is reproduced by its homologue copper ATPase, ATP7A (Menkes disease protein). Interestingly, under basal condition, ectopically expressed mKO-ATP7A shows tight TGN localization (Fig 2G *top panel*) without any localization on endolysosomes (Fig. 2G *bottom panel*). However, mKO-ATP7A exhibited copper responsive exit from the TGN in high copper (Fig. S2C).

To understand the underlying mechanism of unique localization of ATP7B on endolysosomal compartments in hepatic cells in basal copper conditions, we checked if these cells have intrinsically higher copper content compared to non-hepatic cells. Upon measuring intracellular copper, we found that HepG2 contains significantly higher copper as compared to HEK293T and HeLa cells (Fig. S2D). This is an expected observation, as liver serves as the primary detoxifying tissue for all transition metals including copper and hence copper concentration is found to be higher in liver and bile as compared to other tissues, e.g., kidney, stomach, intestine and lung ^37^. Though HEK293 (cell line of kidney origin) and HeLa cells express ATP7B endogenously, these tissues possibly do not face a situation of copper overload in normal day-to-day physiological condition, and hence might not require a constitutive TGN-endolysosomal exchange of copper. Upon quantifying transcript and proteins levels of major candidates of copper homeostasis pathway, i.e., CTR1, ATOX1, and ATP7A, we did not detect any major variation that might be responsible for variable copper loads in these cell lines (Fig. S2E, S2F). However, HEK293T cells exhibited higher levels of ATP7A mRNA and protein that possibly acts as the major copper exporter in these cells.

It is worthwhile to mention that upon prolonged copper chelation in HepG2 cells that resulted in severe copper depletion (Fig.1E), endolysosomal localization of ATP7B persisted (Fig.1F). This chronic chelation of copper, i.e., 10uM BCS for 72h followed by 100uM BCS for 2h resulted in a depletion of ∼80% intracellular copper as compared to non-hepatic cells (Fig.S2D)

From these experiments, and we conclude that copper independent endo-lysosomal localisation is a specialized feature of ATP7B when expressed exclusively in hepatic cell lines. A model in Fig.2H compares cu-independent versus Cu-dependent trafficking of ATP7B in hepatic and non-hepatic cells.

### ATP7B recycles between TGN-proximal lysosomes and the TGN in a copper-independent mode

Upon close examination, we found that the TGN and lysosomes are in close proximity in multiple locations in the perinuclear region of the cell. Interestingly, we found out that an appreciable fraction of ATP7B is localised on these TGN-proximal lysosomes. We observed similar localisation pattern of ATP7B using various TGN and endo-lysosomal marker combination i.e., TGN46-LAMP2 (Fig. 3A), TGN46-LAMP1 (Fig. S3A), Golgin97-LAMP1 (Fig. S3B) and TGN38-LAMP2 (Fig. S3C). ATP7B localizing at TGN46 and late endosome marker Rab7 was significantly diminished as compared to the *bona fide* lysosomal markers (Fig. S3D). Not only in HepG2, but we observed similar phenomenon for another hepatic cell line Huh7 (Fig. S3E). We calculated that under basal copper condition a fraction of ATP7B (5.3±4%, mean±SD) was present where TGN marker TGN46 and endo-lysosomal maker LAMP2 colocalizes.

**Figure 3:**
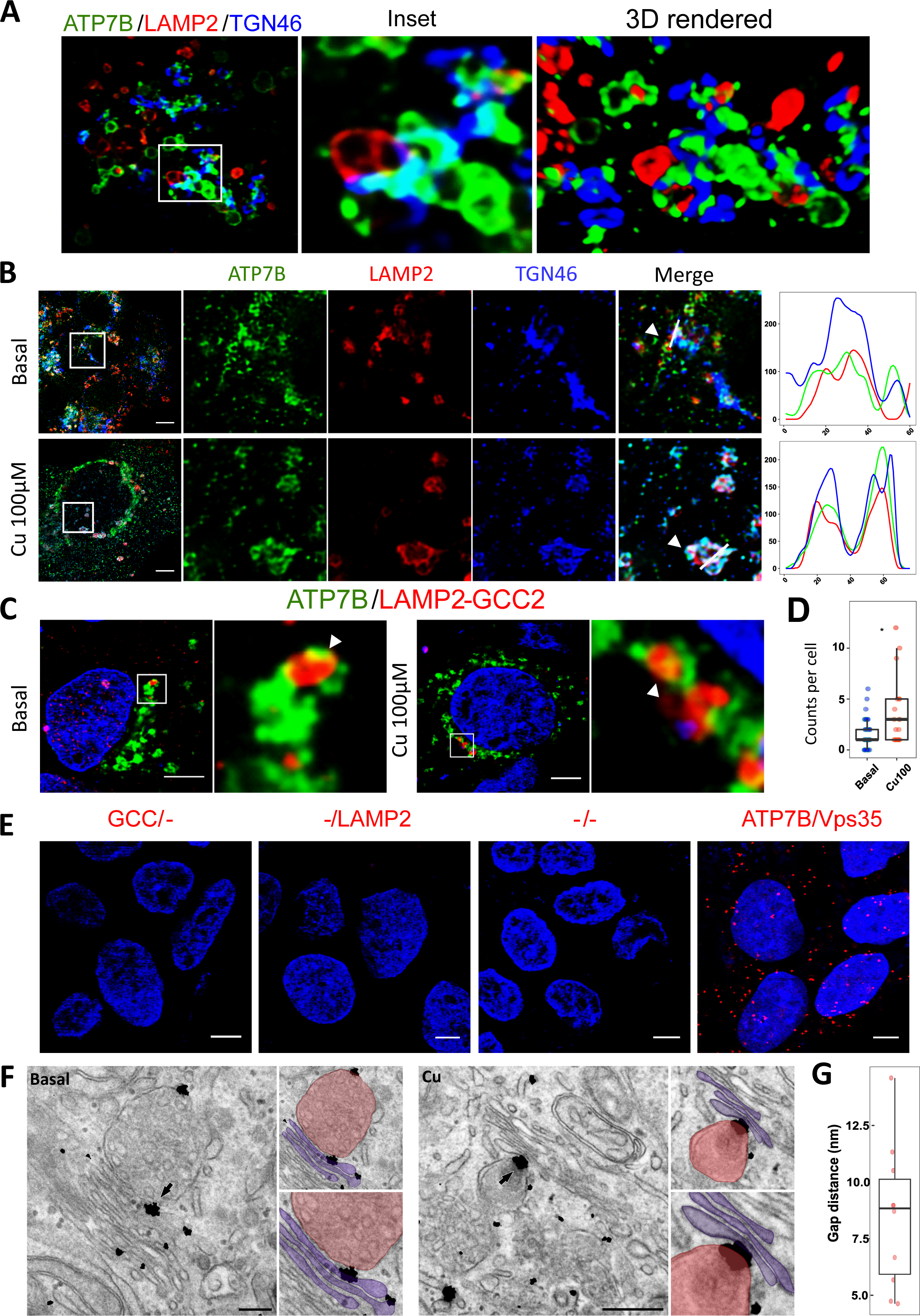
ATP7B localizes at TGN-proximal lysosomes. **(A)** 3D rendered image of ATP7B (green) co-stained with LAMP2 (red) and TGN46 (blue) shows ATP7B localization in TGN46-LAMP2 proximity region in HepG2 cells. **(B)** STED image of ATP7B (green) co-stained with TGN46 (blue) and LAMP2 (red) shown presence of ATP7B in TGN46-LAMP2 colocalizing region, both in basal (top) and Cu (bottom) treated conditions (marked with arrowhead). Marked area in the first panel is enlarged in the next panels. Intensity profile upon the drawn line is represented on the extreme-right panel. **(C)** Proximity ligation assay shows presence of ectopically expressed mEGFP-ATP7B (green) on the GCC2-LAMP2 PLA positive region (red), both in basal (left) and Cu (right) treated conditions (marked with arrowhead). For each condition, marked regions on the left are enlarged in the next. **(D)** Quantitation of PLA positive punctate in basal and copper treated condition is demonstrated by box plot with jitter points. Sample size (N) for (E) are Basal: 42 Cu: 20. **(E)** Immunofluorescence image of negative (only GCC2, only LAMP2, no antibody) and positive control (ATP7B/Vps35) for PLA. **(F)** Immuno-EM of endogenous ATP7B in HepG2 cells shows presence of ATP7B over the TGN-Lysosome proximity sites (marked by arrow) both for the basal (left) and copper treated conditions (right). The enlarged images are presented to the right of each conditions. The Golgi and the lysosomes are colour coded as purple and red respectively. **(G)** The gap distance between Golgi and lysosome at the TGN-lysosome proximity site are represented as box plot with whiskers (N=10). [scale bar for EM: 100 nm, for immunofluorescence: 5µm]

To obtain a higher spatial resolution, we utilised stimulated emission depletion (STED) microscopy to look further into ATP7B distribution at the TGN and the TGN-proximal lysosomes. Our inference of ATP7B localisation on TGN-proximal lysosomes stayed valid (Fig.3B) irrespective of cellular copper condition.

We utilized proximity ligation assay (PLA) to further substantiate our finding that TGN and the TGN-proximal lysosomes constitutively harbouring ATP7B (copper-independent) shares close apposition. In PLA, if two proteins labelled by their respective antibodies are within the proximity range of 40nm, a spot fluorescent emission is detected through in-cell PCR ^38^. We labelled endolysosomes with LAMP2 antibody and TGN with antibody against GCC2. We detected several fluorescent spots when these two antibodies were simultaneously employed. Furthermore, we found some of the PLA signal overlapped with ectopically expressed EGFP-ATP7B (Fig.3C). Interestingly, proximity of lysosomes and TGN increased upon copper treatment as detected by an elevation of PLA signals (Fig.3D). We used ATP7B and VPS35 as the positive control as these two proteins exhibit spatial interaction as shown earlier by our group ^9^ (Fig.3E). No PLA signal was observed in the negative controls (Fig.3E). Overlap of PLA (red) signal with ATP7B (green) confirmed the presence of ATP7B on TGN-proximal lysosomes.

To further substantiate our findings, we performed immuno-EM for endogenous ATP7B in HepG2 cell. Our observation regarding presence of ATP7B on TGN proximal lysosome remained valid (Fig.3F). By measuring the distance between these two organelles, we found it ranged between 5-15 nm (Fig.3G). For the ER-organellar contact sites, the distance between ER and apposing organellar lies between 3-15 nm ^39^. The gap in mitochondria-lysosome contact site was found to be ∼10 nm^40^. There are some other organellar contact sites (e.g. Golgi-Lysosome contact site) where the gap distance is 30 nm ^41–43^. A gap of 10 nm between two organelles can be considered as a potential organellar contact sites ^41^. Interestingly, in basal copper conditions, immune-EM revealed the presence of ATP7B at sites of proximity between TGN and Multivesicular bodies (MVB) marked by inclusion structures within them (Fig.3F, *left panel*). ATP7B was found in TGN-lysosome proximity sites in conditions of high copper (Fig.3F, *right panel*). Our findings show that ATP7B lies in close proximity sites between TGN and organelles marked by LAMP2. With the present set of data, it is difficult to ascertain the maturation stage(s) of lysosome at these proximity sites that harbors ATP7B. To summarize, we hypothesise the existence of contact sites between TGN and lysosome (or endolysosomal compartments and MVBs) and presence of a fraction of ATP7B therein. However, all such proximity sites did not harbour ATP7B.

Proximity of organelles and their biological relevance have been recently studies especially between TGN-endoplasmic reticulum (ER) and lysosome-ER. Using colocalization studies using organelle-specific markers, i.e., Calnexin and TGN46 for ER-TGN and LAMP2 and Calnexin for lysosome-ER and we excluded the chances of ATP7B being present on these proximity sites (Fig.S3F, S3G).

LAMP2 is a heavily glycosylated recycling protein that is also detected at the trans-Golgi network in the late secretory pathway ^44(p41)^. To inspect if the TGN-lysosomal overlap is formed by ‘TGN-localised’ or ‘TGN-exiting’ LAMP2 in the late secretory pathway or this lysosome is a part of the maturing endocytic pathway, we performed a dextran uptake assay. Dextran is up taken by cell via endocytosis and then localises to acidic compartment by the process of endosomal maturation. Therefore, any overlap between TGN and Dextran can be accepted as a *bona fide* TGN-lysosome proximity site. We observed colocalization between TGN marker S-galactosyl transferase and acidic compartments (dextran positive) and ATP7B in basal copper condition (Fig.4A). Hence, we can confirm that a fraction of ATP7B constitutively localizes at those TGN-lysosome proximity regions.

**Figure 4:**
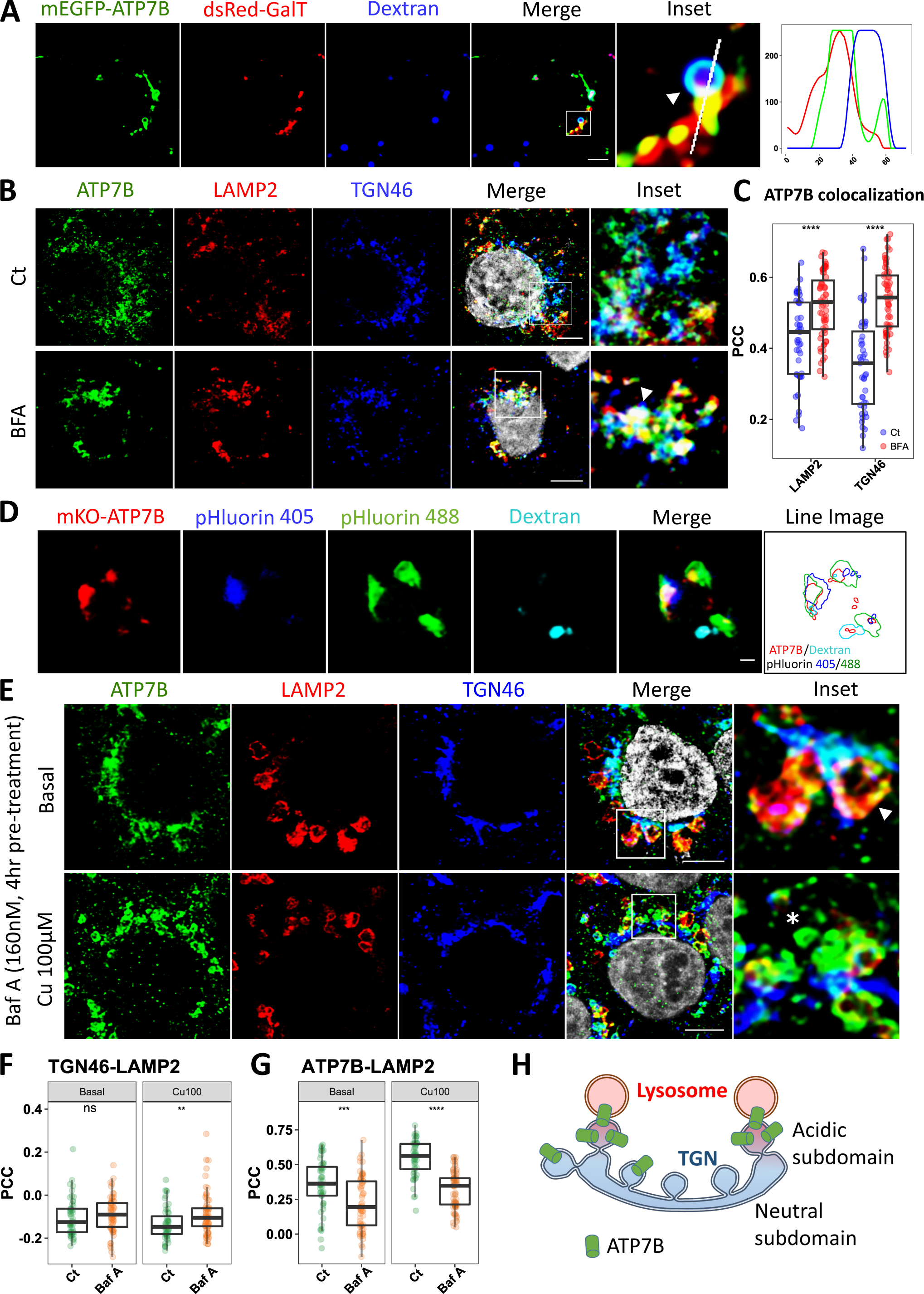
ATP7B localization at the TGN and lysosomes and at TGN-lysosomes colocalized compartments upon TGN disruption. **(A)** Ectopically expressed mEGFP-ATP7B (green) is co-expressed with dsRed-GalT (red). Acidic compartments are marked with Dextran (blue). Localization of ATP7B in the region where Golgi (red) and acidic compartment (blue) colocalizes shows that ATP7B is present on ‘true’ TGN-proximal lysosome (marked with arrowhead). Intensity profile upon the drawn line is represented on the extreme-right panel. **(B)** Immunofluorescence image of ATP7B (green) in HepG2 cell line (control, *top panel* or Brefeldin-A treated, *bottom panel*) co-stained with LAMP2 (red) and TGN46 (blue) shows that Brefeldin-A treatment results in colocalization of TGN46 and LAMP2 and presence of ATP7B therein, marked with arrow (bottom panel). **(C)** The PCC between ATP7B-TGN46 and ATP7B-LAMP2 in basal condition in control, ct (green) and BFA (orange) treated conditions are demonstrated by box plot with jitter points. Sample size (N) are ct: 43, BFA: 63. **(D)** Subdomain of TGN based on pH is marked by ratiometric pHluorin tagged TGN38 in blue (low acidity) and green (acidic). [Scale bar: 1µm] Line image shows the relative position of all the proteins. **(E)** Immunofluorescence of ATP7B (green) co-stained with TGN46 (blue) and LAMP2 (red) shows altered trafficking of ATP7B in high Bafilomycin-A_1_ pre-treatment (160nM for 4h), both in basal (top) and Cu (bottom) treated conditions (marked with arrowhead). Marked area in the first panel is enlarged in the next panels. The arrowhead shows colocalization between ATP7B-LAMP2 (top panel), the asterisk shows mislocalized ATP7B near TGN46 (bottom panel). The PCC between LAMP2-TGN46 **(F)** and ATP7B-LAMP2 **(G)** in basal and copper treated conditions in control, ct (green) and Bafilomycin-A1 (orange) pre-treated conditions are demonstrated by box plot with jitter points. Sample size (N) are for ct: Basal: 52, Cu: 52; for Baf-A treated: Basal: 62, Cu: 71. **(H)** Model shown presence of ATP7B in the acidic sub-domain of TGN in close proximity of lysosome. [The scale bar for rest of the figures: 5µm.]

Lysosomes are dynamic organelles which sometimes lie in close apposition with different other organelles. Using proximity ligation Hao et al. has shown that lysosome possibly makes contacts with cis-Golgi forming unique Golgi-Lysosome contact sites (GLCS) that is important for mTOR activation in cell ^42^. We investigated if GLCS resides at the same zone where TGN-lysosomes share high proximity. Interestingly, we have found that GLCS formed by cis-Golgi marker GM130 and endolysosomal marker LAMP1 are quite distinct from TGN46-LAMP1 proximity regions (Fig.S4A). Moreover, ATP7B was not present on GLCS (Fig.S4B). This data suggests our hypothesised TGN-Lysosome/MVB contact site is unique and different than previously described GLCS.

We argued that if TGN and endolysosomes share high proximity and ATP7B is constitutively exchanged between these two organelles, then disrupting the Golgi should lead to rapid exchange between these two organelles leading to increase in colocalization between endolysosomal and TGN markers. Interestingly, we observed that disrupting the Golgi using Brefeldin-A resulted in formation of punctate compartments that was positive for both LAMP2 as well as TGN46 as evident from significantly increased colocalization between TGN46 and LAMP2 (Fig.S4C). As hypothesized, ATP7B was found in these colocalized TGN-endolysosome domains upon Golgi disruption (Fig.4B, 4C). This observation suggests the presence of a possibly direct and constitutive ATP7B-containing membrane exchange phenomenon between the two organelles.

### ATP7B localizes at the region of the trans-Golgi network that exhibits lower pH

TGN and the proximal lysosomes constitutively exchange ATP7B. The TGN and lysosome lumen exhibits different pH, lysosomal lumen being more acidic as compared to lumen of Golgi stacks. A large amount of the ATP7B was found to be present in a milieu around the TGN and the proximal lysosomes (0.5-1.5µm). Copper ATPases are known to require slightly acidic environment for its efficient copper pumping activity ^45,46^. Since copper pumping activity of ATP7B is pH dependent, we tested if ATP7B is preferentially present on any subdomain of TGN based on pH. We used pH-sensitive pHluorin tag attached to the C-terminal of TGN38 which anchors it to the TGN membrane. The sensor, pHluorin is a modified GFP that has two pH-dependent excitations. At neutral pH, it is excitable at 395nm where at acidic pH, it gets excited at 480nm. With decreasing pH, the emission at 480nm increase as well as ratio of the two emissions Em_480_/Em_395_ ^47^. So, from two excitations, we can gather a qualitative idea of the luminal pH gradient (if any) and correlated preferential localization of ATP7B within the TGN. Upon expressing the pHluorin plasmid in HepG2 cells, we found that ATP7B is preferentially present on the acidic sub-domain of TGN as visualized by its higher colocalization with pHluorin when excited by the wavelength of 480nm as compared to excitation at 395nm (Fig.4D). We also observed that the acidic subdomain of TGN shares close proximity with lysosomes marked by dextran (Fig.4D). We can infer that the low pH organellar subdomains act as the sites of exchange of ATP7B between the TGN and TGN-proximal lysosomes.

We further check the luminal pH of TGN plays a role on distribution of ATP7B at the TGN and TGN-proximal endolysosomes. It has been demonstrated by Levic and coauthors. that optimum pH maintenance by the V-ATPase pump is required for sorting and trafficking of apical membrane proteins in zebrafish intestine ^48^. Previous studies have demonstrated that disrupting lysosomal acidification does not affect copper-mediated lysosomal localization of ATP7B ^9,49^. Using 40nM of Bafilomycin A1 for 30min pre-treatment, which is enough to disrupt lysosomal acidity (as shown by rapid decrease in Lysotracker intensity in flow cytometry measurement, Fig. S4D) did not affect ATP7B localization appreciably either in TGN or Lysosome in both basal and copper treated condition (Fig.S4E, S4F). Interestingly, pre-treatment with a higher concentration of Bafilomycin A for a longer period (160nM for 4h) affected ATP7B localization significantly (Fig.4E). Though colocalization between TGN46 and LAMP2 increased slightly (Fig.4F), ATP7B localization at LAMP2, both in basal and Copper treatments decreased significantly (Fig. 4G). Previous studies have shown that altering Golgi acidification blocks protein trafficking from Golgi ^50^. Furthermore, in ‘high’ Bafilomycin A treatment, in high copper, ATP7B was found to exit TGN only to create blob-like structure attached to it, similar to what was reported by Lalioti and co-authors ^51^. Upon Bafilomycin A treatment, larger LAMP2 positive endolysosomes were formed at the perinuclear position (Fig.4E). Hence, not only TGN-Lysosome proximity, but TGN lumen pH is also important for proper trafficking of ATP7B. The model of pH dependence of ATP7B trafficking is illustrated in Fig.4H.

### TGN-proximal lysosomes serve as a hub for endo-lysosomal sorting of ATP7B

ATP7B traffics to endolysosomes from TGN upon elevated intracellular copper levels. It is well documented that ATP7B is directly transported to endo-lysosomal compartment (late secretory pathway) and not by endocytic pathway of lysosomal maturation via plasma membrane ^6^. To circumvent copper-induced toxicity, this type of direct transport ensures faster response in elevated copper condition. A recent study has shown that TGN associated lysosome acts as a hub for secretory, endosomal and autophagic route in drosophila ^52^. Interestingly, we found that increase in TGN46-LAMP2 colocalization is positively correlated with copper-induced endolysosomal localisation of ATP7B that increases with increasing copper (Fig.5A, 5B). We further explored if these TGN-proximal lysosomes serve as a ‘*distribution hub’* for endolysosomal dispersal (trafficking) of ATP7B in elevated copper. First, we conducted time-lapse imaging of ATP7B trafficking during copper treated condition. Interestingly, we have found that dextran positive acidic compartments to be reaching in contact with the TGN upon 100µM copper treatment. ATP7B signal was found to increasing in the dextran positive compartment and simultaneously decreasing at the TGN which suggests possible ATP7B exchange between these two compartments (Fig 5C, Video 1). These findings reveal that high copper induces localization of endolysosomal compartments at the TGN where ATP7B exchange occurs between the TGN (donor compartment) and the lysosomes (acceptor compartment).

**Figure 5:**
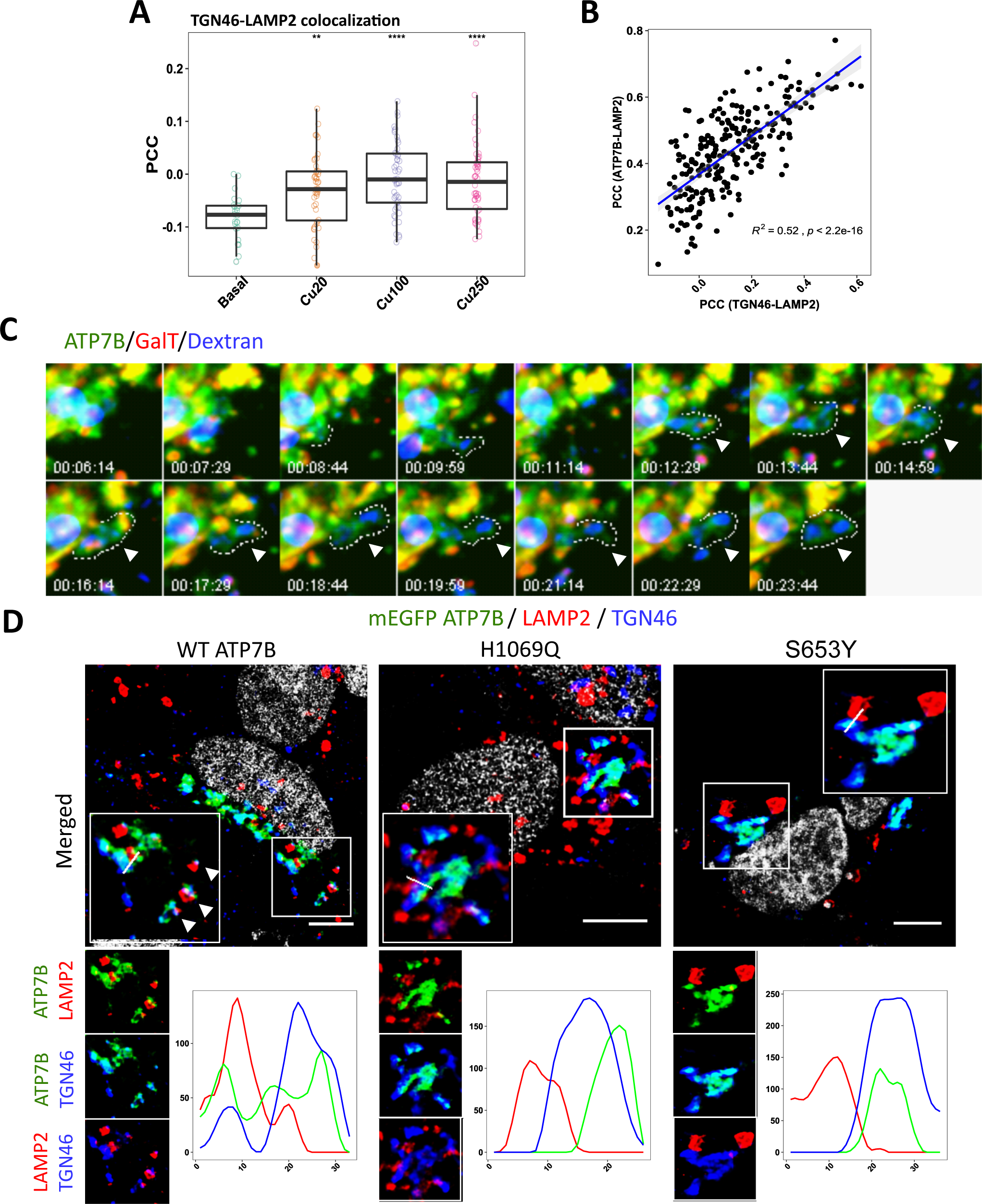
TGN-proximal lysosome plays a role in the endo-lysosomal sorting of ATP7B. **(A)** PCC between TGN46-LAMP2 shows increased colocalization upon copper treatment. The data is represented as box plot with jitter points. Sample size (N) for Basal: 24, Cu20: 44, Cu100: 49, Cu250: 50. **(B)** Regression model shows a positive relationship between TGN46-LAMP2 and ATP7B-LAMP2 PCC values. **(C)** Time lapse imaging of ectopically expressed mEGFP-ATP7B (green) co-transfected with dsRed-GalT (red). Acidic compartments are marked with Dextran (blue). Arrowhead shows pinching off of ATP7B from Golgi to a Dextran positive vesicle. Time scales are in minutes. **(D)** Ectopically expressed mEGFP-ATP7B (wild-type, H1069Q, and S653Y) co-stained with LAMP2 (red) and TGN46 (blue) shows presence of Golgi exiting ATP7B in TGN46-LAMP2 colocalizing region (marked with arrow-head). Marked area in the first panel is enlarged as well as two protein co-staining is shown in bottom-left panel. Intensity profile upon the drawn line is presented on the bottom-right panel. [Scale bar: 5µm]

Since no biochemical method is known to disrupt or manipulate for TGN-proximal lysosomes, we employed indirect methods to alter ATP7B localization at these sites. More than 400 WD causing mutations are known for the ATP7B protein. Many of them results in defective trafficking of the protein ^53^. H1069Q, the most prevalent mutations reported in Caucasian population exhibits compromised copper-induced Golgi exit of the protein ^54^. Another WD mutation S653Y though capable of copper transport at the Golgi, shows complete non-trafficking phenotype ^26^. We tested if these mutants differ as compared to the wt-ATP7B with respect to their presence on the TGN-proximal lysosomes. Upon expressing EGFP-tagged mutant ATP7B and the wt-ATP7B, we found that Golgi retained mutants (H1069Q and S653Y) show much reduced TGN-proximal lysosomal localisation than wildtype ATP7B (Fig.5D). This observation indicates that ‘trafficking-ready’ ATP7B resides at these TGN-proximal vesicles that possibly serves as the exit point for copper-induced trafficking.

### Lysosomal positioning regulates ATP7B abundance at the TGN-proximal lysosomes

Lysosomal positioning is one of the major indicators of cellular health. Depending upon nutrient status of the cell, lysosomes could be either attain perinuclear or peripheral localization in a mammalian cell ^14,15,55,56^. In addition to that, endolysosomes that are at the proximity to TGN are essentially perinuclear lysosome. So, we wanted to ask if lysosomal positioning at these TGN-proximal sites or at the cell periphery have any effect on localisation of ATP7B. We overexpressed lysosomal position-modulating proteins to enrich either perinuclear or peripheral lysosomes.

RUFY3 is an important modulator for retrograde trafficking of lysosome. It is a member of a small group of proteins called RUFY or “<u>RU</u>N and <u>FY</u>VE domain-containing protein” that acts as adaptor to small G-proteins specifically ^57^. RUFY3 is an effector of Arl8b that promotes the formation JIP4-dynein-dynactin complex and thus modulates retrograde trafficking of Lysosome. Overexpressing RUFY3 promotes perinuclear localisation of lysosome^58^. On the other hand, overexpressing Arl8b results in increased lysosomal localisation towards cell periphery. Arl8b is an Arf-like small GTPase that plays a crucial role in regulating lysosomal positioning. Like other GTPases, upon GTP bound condition, the active form of Arl8b localises on lysosomes and promotes anterograde trafficking of lysosome towards periphery ^58–62^.

In Arl8b overexpressed cell, we did not observe any change in endolysosomal localisation of ATP7B in basal or copper treated cells (Fig.6A, 6B). However, there was a significant decrease in the abundance of TGN-proximal lysosomes as determined by calculating the signal overlap between LAMP2 and TGN46 (Fig.6C). In cells overexpressing RUFY3, we observed increased endo-lysosomal localization of ATP7B and decreased TGN localisation than un-transfected cells (control) at basal and copper treated condition (Fig.6D, 6E). However, though we observed only a slight increase in TGN-proximal lysosomes, the difference was non-significant (Fig.6F). Cells overexpressing Arl8b and RUFY3 (pseudo-coloured in purple) have been shown in Fig.S5A.

**Figure 6:**
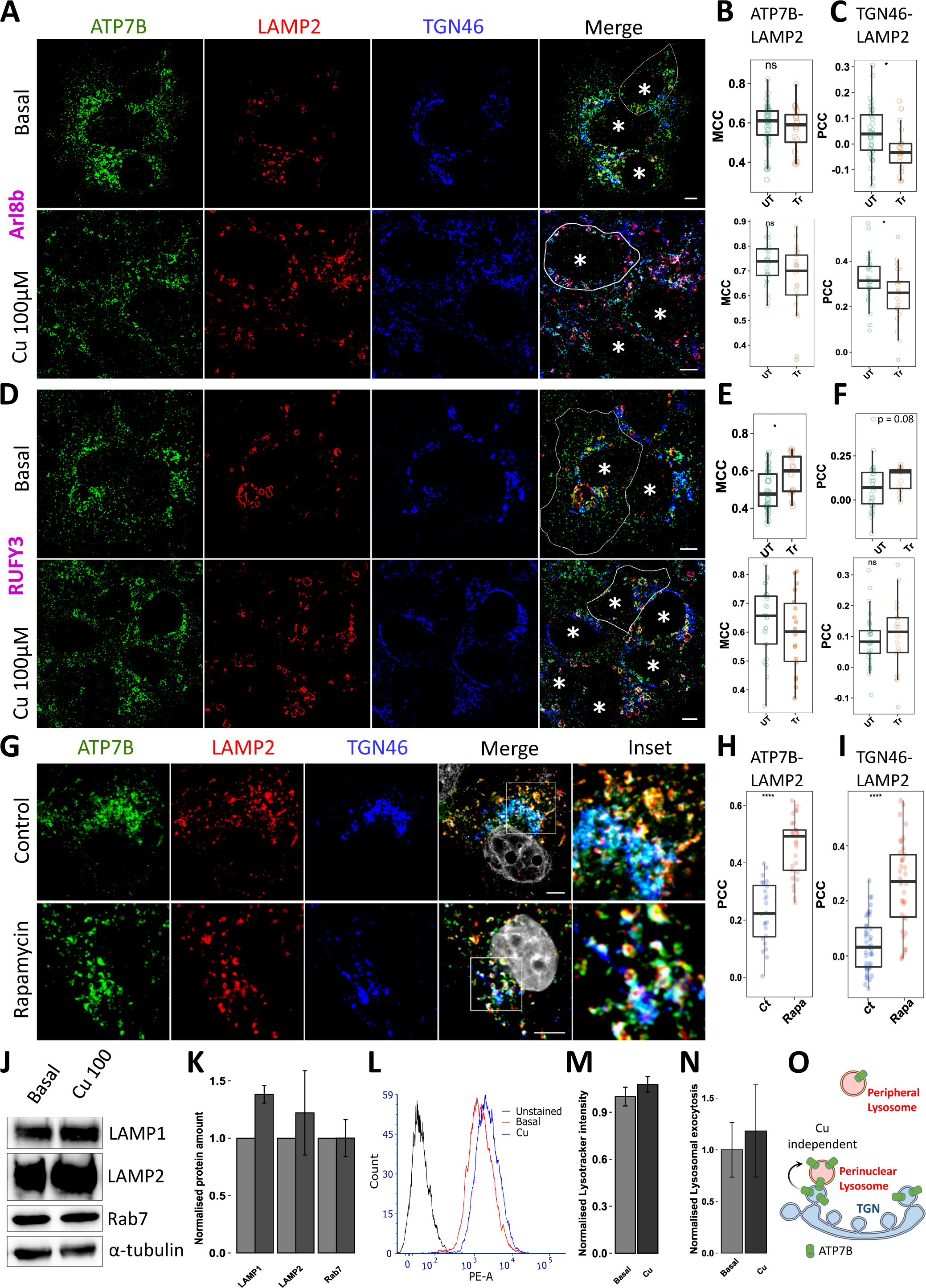
Lysosomal positioning regulates ATP7B abundance on the TGN-proximal lysosome. **(A)** Immunostaining of ATP7B (green) co-stained with LAMP2 (red) and TGN46 (blue) in HepG2 transfected (Tr) with GFP-Arl8b in un-transfected cells (UT) in basal and copper treated condition. Boundaries of transfected cells are marked with white line. Asterisk marks the nucleus of the cells. The Fraction of ATP7B over LAMP2 **(B)** and PCC between TGN46-LAMP2 **(C)** in transfected and un-transfected cells are demonstrated by box plot with jitter points. Sample size (N) for Basal are Tr: 20, UT: 38; for Cu treated are Tr: 21, UT: 29. **(D)** Immunostaining of ATP7B (green) co-stained with LAMP2 (red) and TGN46 (blue) in HepG2 transfected (Tr) with GFP-RUFY3 in un-transfected cells (UT) in basal and copper treated condition. Boundaries of transfected cells are marked with white line. Asterisk marks the nucleus of the cells. The Fraction of ATP7B over LAMP2 **(E)** and PCC between TGN46-LAMP2 **(F)** in transfected and un-transfected cells are demonstrated by box plot with jitter points. Sample size Basal are Tr: 14, UT: 42; for Cu treated are Tr: 18, UT: 29. **(G)** Immunofluorescence image of ATP7B (green) in HepG2 cell line co-stained with LAMP2 (red) and TGN46 (blue) shows that Rapamycin treatment results in increased ATP7B localization in endolysosomes. The PCC between **(H)** ATP7B-LAMP2 and **(I)** TGN46-LAMP2 in basal and copper conditions in ct (green) and Rapamycin (orange) treated conditions are demonstrated by box plot with jitter points. Sample size (N) are ct: 43, Rapa: 37. **(J)** Immunoblotting and **(K)** subsequent quantitation of LAMP1, LAMP2 and Rab7 in basal and copper treated condition shows lysosomal biogenesis upon 2h copper treatment. Loading control: α-tubulin. Expression level is normalized against loading control as well as basal. Data is demonstrated as bar-plot (mean ± SD). **(L), (M)** Flow cytometry analysis shows increase in Lysotracker intensity upon copper treatment. Data is normalized against basal and demonstrated as bar-plot (mean ± SD). **(N)** Lysosomal exocytosis increases in response to copper treatment. Data is normalized against basal and demonstrated as bar-plot (mean ± SD). **(O)** Model shows copper independent trafficking of ATP7B at the peri-nuclear lysosome. [Scale bar: 5µm]

We further modulated mTOR to manipulate the distribution of peripheral and perinuclear lysosomes. mTOR (<u>m</u>ammalian <u>T</u>arget <u>O</u>f <u>R</u>apamycin) is a serine/threonine kinase that, upon activation, is present on lysosomal membrane ^63^. Several studies have elucidated the regulatory mechanism of lysosomal recruitment and activation by mTORC1 that follows the established ‘canonical’ as well as the more recently described ‘non-canonical’ pathways ^14,64,65^. The mTOR protein forms complexes (mTORC1 and mTORC2) by regulating lysosomal functionality and localization affects a spectrum of cellular processes ^16,17,66^. In elevated copper condition, mTOR is known to get inhibited by some unknown mechanism ^29^. Cu mediated inactivation of mTOR leads to upregulation of TFEB which in turn upregulates lysosomal biogenesis to exocytose extra copper from cell ^7,67^. Further, inhibiting mTOR using rapamycin also results in perinuclear localisation of lysosomes. So, we wanted to create a similar Cu-stressed environment inside cell in absence of copper. Interestingly, in rapamycin treated cells, we observed elevated lysosomal localisation of ATP7B than TGN localisation irrespective of cellular copper condition (Fig.6G, 6H). Moreover, colocalization between TGN46 and LAMP2 was also rises indicating an increase in TGN-proximal lysosomes (Fig.6I). This finding was further validated with proximity ligation assay where we recorded an increase in LAMP2-GCC2 PLA positive spots in copper compared to basal condition that further increase upon Rapamycin treatment (Fig.S5B, S5C). Not only an increase in TGN-lysosomal localization, copper treatment leads to increased lysosomal biogenesis which was measured by immunoblotting LAMP1 and LAMP2. We did not observe any change in late endosomal marker Rab7 level (Fig.6J, 6K). In addition, we also observed slight increase in lysotracker intensity labelling the lysosomes in flow cytometry analysis of cells in copper treated condition (Fig.6L, 6M). We also observed an increase in lysosomal exocytosis upon copper treatment as determined by hexosaminidase activity (Fig.6N) which corroborates with previous findings by Polishchuk and co-workers ^6^.

To summarize, increasing the abundance of perinuclear lysosomes augments endolysosomal localization of ATP7B that happens via exchange of the protein between the TGN and its proximal lysosomes (illustrated as a model in Fig.6O). Also, to efficiently export excess copper from the cell by lysosomal exocytosis, the cell employs increased lysosomal biogenesis in elevated copper.

### ATP7B recycles back to TGN-proximal lysosomes upon copper chelation

In elevated copper condition, ATP7B exports excess copper via lysosomal exocytosis. Subsequently, upon copper chelation, ATP7B recycles from endolysosome in a retromer regulated pathway ^9^. So next, we asked whether the vesicularized (trafficked) ATP7B, upon copper-chelation completely recycles back to the TGN. For studying the retrograde trafficking of ATP7B, we treated cells with 250μM of copper for 2 h and then chelated with 100μM of copper chelator, BCS and recorded its trafficking using time lapse imaging. Interestingly, while tracking the retrograde pathway using live imaging, we found that acidic compartments marked by dextran carrying ATP7B returns from peripheral location to perinuclear location but the endolysosomal pool of ATP7B does not relocalize to the TGN. Rather, it remained localised in acidic compartments that are juxtaposed to the TGN (Fig.7A and video 2). Our observation holds true even if we continued imaging for an extended time. To avoid skewing the abundance of ATP7B in TGN pool due to *de novo* synthesis of the protein, we halted protein synthesis using cycloheximide (CHX) during the entire time period of the experiment. We found that, in cycloheximide treated condition, even after copper chelation, the TGN pool and the total endosomal pool of ATP7B remained similar as copper treated condition. Interestingly, upon triggering the retrograde pathway of ATP7B with the copper chelator BCS for 30 mins or continued for 2h, ATP7B was detected in peri-Golgi lysosomes with a fraction of it localizing at very close proximity to the TGN (shown by white arrows in Fig. 7B, bottom panel of CHX treatment, magnified insert and Fig S5D). On the contrary, in the control set (not treated with cycloheximide), the TGN localisation of ATP7B increases upon copper chelation which suggests that the TGN localised ATP7B pool is due to newly synthesized ATP7B during the entire experimental process (Fig.7B, 7C, 7D). In CHX treated cells, colocalization between ATP7B and LAMP2 and the fraction of ATP7B on LAMP2-labelled endolysosomes increases with copper treatment and stays high even after activating the retrograde pathway, compared to control cells (Fig.7C, S5E *top panel*). Understandably, a reverse phenomenon was observed in case of ATP7B and TGN46 colocalization and the fraction of ATP7B at the TGN46-labelled trans-Golgi network (Fig.7D, S5E *bottom panel*). The comprehensive comparative analysis and box-plot of colocalization of ATP7B with LAMP2 and TGN46 between CHX treated and control cells the four copper conditions have been illustrated in Fig.S5F. This finding suggests that endolysosomal fraction of ATP7B remains at the endolysosomes after activating the retrograde pathway. The retrograde pathway of ATP7B is possibly an ‘*incomplete recycling pathway*’ ensuring that recycled ATP7B is not reinstated back to the TGN but stays in the endolysosomes closer to the peri-Golgi region of the cell. This provision might be beneficial for a quick response to elevated copper levels when ATP7B needs to conduct copper export via lysosomal exocytosis. A model illustrates the retrieval of ATP7B from peripheral lysosomes to TGN-proximal lysosomes in Fig.7E. To gather a higher resolution of the phenomenon that the retrograde trafficking of ATP7B culminates at the lysosomes, we conducted immune-EM. Remarkably, as recorded with light microscopy we observed that indeed ATP7B exhibits localization at the lysosomes even after the retrograde trafficking was triggered by BCS, post copper-mediated vesicularization of the protein (Fig.8A)

**Figure 7:**
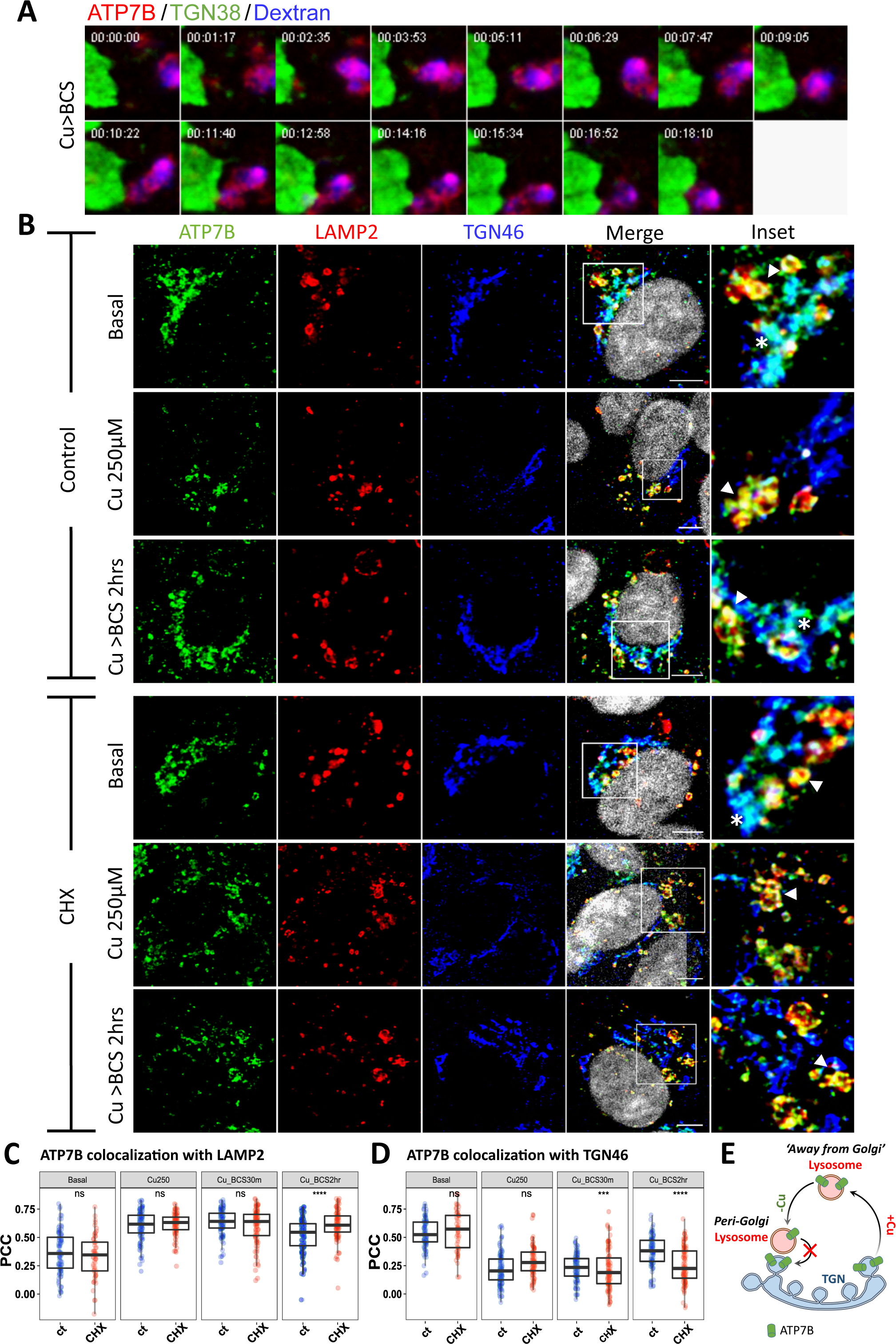
Vesicularized ATP7B recycles back to the TGN-proximal lysosome upon copper chelation. **(A)** Time lapse imaging of ectopically expressed mKO-ATP7B (red) co-transfected with pHluorin tagged TGN38 (green). Acidic compartments are marked with Dextran (blue). Time scales are in minutes. **(B)** Immunostaining of ATP7B (green) co-stained with LAMP2 (red) and TGN46 (blue) in HepG2 in control and CHX treated conditions in different copper conditions i.e., basal, Cu250 and Cu250>BCS, 2hr. The arrowhead shows colocalization between ATP7B-LAMP2, the asterisk shows colocalization between ATP7B-TGN46. The PCC between ATP7B-LAMP2 **(C)** and ATP7B-TGN46 **(D)** in control (ct) and CHX treated cells under different copper conditions are demonstrated by box plot with jitter points. Sample size (N) for (C) and (D) in basal condition are basal: 94, Cu250: 108, Cu_BCS30m: 104, Cu_BCS2hr: 99. Sample size for CHX treated condition are basal: 57, Cu250: 84, Cu_BCS30m: 100, Cu_BCS2hr: 103. **(E)** Model shows that ATP7B does not recycle back to the TGN upon copper chelation. [Scale bar: 5µm.]

**Fig 8:**
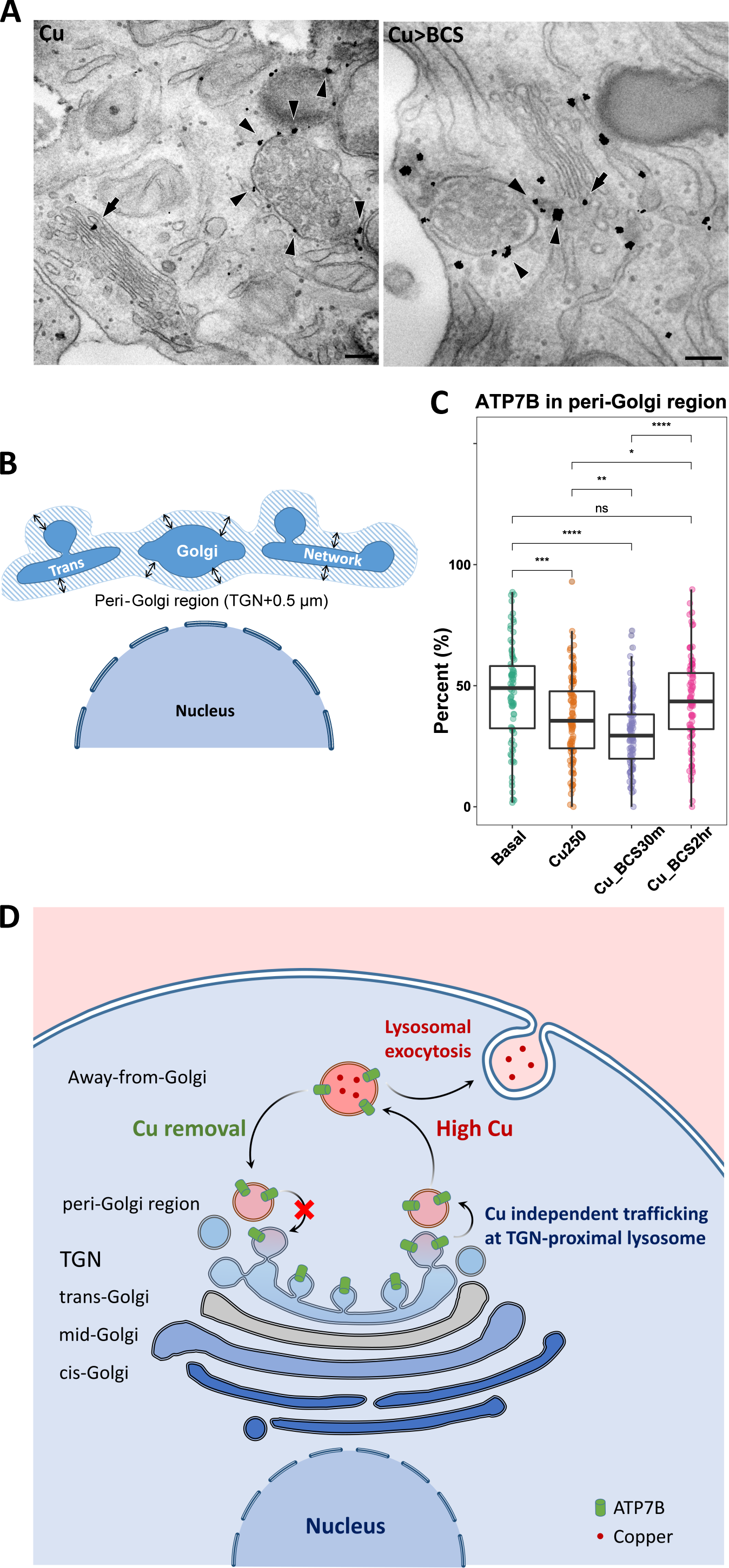
Vesicularized ATP7B recycles back to peri-Golgi lysosomes and TGN-proximal lysosome upon copper depletion. **(A)** Immuno-EM of endogenous ATP7B shows redistribution of ATP7B to Lysosome upon copper treatment (left panel, marked by arrowhead). Upon copper chelation, the ATP7B was detected on the peri-Golgi region (right panel, marked by arrowhead). Golgi Localized ATP7B are marked with arrow. **(B)** Model representing peri-Golgi region that harbors the peri-Golgi lysosomes **(C)** Quantitation of Localization of ATP7B in peri-Golgi region upon copper treatment and subsequent copper chelation. Sample size (N) in basal condition are basal: 80, Cu250: 100, Cu_BCS30m: 95, Cu_BCS2hr: 86. **(D)** Model illustrating the copper-induced trafficking of ATP7B between perinuclear and peripheral lysosomes and its localization at the TGN-proximal lysosome. [scale bar for EM: 100 nm]

To set as clear cut-off distance from the nucleus that will demarcate perinuclear vs peripheral lysosomes is not possible as (a) there always exists a fraction of lysosomes that are either traveling in the anterograde or the retrograde pathway but are yet to reach the point of final destination at the periphery or the perinucleus respectively and (b) the nucleus/cell ratio is large in HepG2 cells and therefore making it difficult to clearly distinguish between the two pools of lysosomes. Since ATP7B traffics between TGN and lysosomes, it is perhaps more apt to use a Golgi centric approach instead of nuclear-centric approach for analysis of ATP7B positioning in our study. So, for the purpose of quantitation and nomenclature, we used peri-Golgi and away-from-Golgi to name two pools of ATP7B. Anything within the range of 0.5µm from the nearest TGN boundary was considered as peri-Golgi and the rest of ATP7B was pooled as away-from-Golgi (Fig.8B). This is because at basal condition most of ATP7B was found at a distance within 0.5µM from the TGN boundary. This pool of ATP7B at peri-Golgi lysosomes at four different copper conditions are depicted in Fig.8C.

To summarize our findings, in hepatocytes, ATP7B exhibits a constitutive localization in the TGN and TGN-proximal lysosomes. These post-TGN lysosomal compartments may serve as the sites where ATP7B accepts copper from ATOX1. Upon elevated copper, lysosomal trafficking from TGN-proximal to peripheral zone leads to higher vesicularization of ATP7B that can now export copper via lysosomal exocytosis. Upon copper chelation, the protein recycles back to the TGN-proximal lysosomes, but not to the TGN, to reinitiate the next round of copper transport. The present study leads us to further hypothesize that constitutive presence of ATP7B on TGN-proximal lysosomes saves the cell of energy expenditure of vesicular budding and fission from the TGN at every cycle of copper export by the Cu-ATPase.

## Discussion

The Wilson disease protein, ATP7B maintains physiological levels of copper in the hepatocytes. It has been well documented that ATP7B traffics in vesicles upon copper treatment in all in vitro models of hepatocyte. At the compartments ATP7B utilizes lysosomal exocytosis to export out the excess copper ^6^. Vesicularized ATP7B exhibits retrograde trafficking upon copper chelation. Previously we have shown that the retromer complex regulates recycling of the protein from the endolysosomal vesicles ^9^. In the present study we described a pool of lysosomes that lies in close proximity to the TGN and that the copper transporting ATPase, the Wilson disease protein, ATP7B localizes at these TGN-proximal lysosomes.

We further asked, why does ATP7B localize at TGN-proximal lysosomes? We believe that these sites act as copper dependent as well as copper-independent TGN exit sites for ATP7B. These sites may be the initial points of the late secretory pathway for ATP7B where the ATPase pumps/transports copper even in the absence of a receptor protein e.g., Ceruloplasmin ^45^. Our hypothesis is further supported by lower pH around these zones where these two organelles share high proximity. Lower pH acts as sites of copper release from the transmembrane domains of ATP7B that harbours the conserved copper-sequestering CPC motif as shown previously by Barry et al ^1,6,45,46^.

We observed that this interaction between the TGN and the TGN-proximal lysosome is very transient. Luzio et al mentioned in the review ^68^ that transport through early endosomes to late endosomes that involves delivery of lysosomal hydrolases occurs through ‘kiss-and run’ events as well as complete fusions between late endosomes and lysosomes. We hypothesize that in case of TGN to lysosome transport as observed in case of ATP7B, there will not be a complete fusion between the two compartments because of major differences in their luminal pH. Also, exchange of lysosomal enzymes upon complete fusion between the two organelles will be unfavorable for normal cell physiology.

In the last few years, organellar interfaces, e.g., ER-endosome contact sites have been shown to be modulators of multiple cellular physiology particularly, lipid transfer or recruitment of various complexes and enzymes at ER-endosome. These contact sites play an active role in endosome sorting, scission, and maturation ^69–73^. Lysosome is a dynamic organelle which is known to make contact sites with different other organelles ^40,42,69,74^. These organelles contact sites provides unique environment inside cell, key for vital physiological functions of these organelles e.g., inter-luminal exchanges. Recently discovered Golgi-Lysosome contact sites (GLCS) has been shown to be crucial in regulating mTORC activity ^42^. In the immuno-EM images (Fig.3F), we noticed clusters of ATP7B localized at sites where TGN and the lysosomal membrane shares close apposition. We hypothesize that these zones might represent a novel TGN-lysosome/MVB contact sites, where extensive exchange of ATP7B between the TGN and endo-lysosomes might occur. These sites might be of a great interest in understanding Golgi-sorting and trafficking of protein from TGN. Since organellar proximity is not the only determining factor for contact sites, further in-depth study to confirm their presence and functionality is warranted. Since LAMP2 labels different types of lysosomes and also the various stages of their maturation, we were unable to establish whether a specific class of lysosome is associated with these proximity sites or this was a general phenomenon with all LAMP2-associated endolysosomal and MVB compartments.

We found that upon activation of the retrograde pathway, ATP7B recycles to the perinuclear zone of the cells and resides at the TGN-proximal lysosomes in contrast to its complete relocalization at the TGN, described previously ^9,75(p1)^. We argued that recycling back from the lysosomes to the TGN is an energetically unfavourable process for the ATPase. As for each consecutive copper export cycles, ATP7B need to bud out in vesicles from the TGN, with subsequent maturation of the vesicles from early secretory vesicles to mature lysosomes where ATP7B will export copper in its lumen. To induce the retrograde trafficking pathway of ATP7B, we depleted cellular copper using extracellular copper chelator, BCS as well as intracellular copper chelator, TTM. We observed that a major pool of ATP7B recycles back not to the TGN but to lysosomes that localize at close apposition with the TGN. To understand the dynamics of ATP7B at the TGN-proximal lysosomes, we manipulated the lysosomal localization directly by overexpressing proteins that either triggers lysosomal trafficking from the perinuclear zone to the cell periphery or *vice versa*. Upon increasing the abundance of TGN-proximal lysosomes, we found a higher colocalization coefficient of ATP7B and LAMP2. This points towards the possible copper-independent ‘*anterograde movement*’ of ATP7B from TGN to endolysosomes via these compartments. Elevated copper not only facilitates this process but also triggers lysosomal biogenesis to efficiently export copper and releases the cell of the copper overload.

By the virtue of close proximity of ATP7B containing lysosomes to the TGN, also places them closer to the MTOC that might facilitate copper export in the bile as the hepatocyte polarizes ^76^. Localization of ATP7B on TGN-proximal lysosomes might be beneficial for copper export. It is interesting to note that perinuclear lysosomes as compared to the peripheral ones are primarily involved in nutrient recycling in cells ^14,77^. It is highly probable that the primary function of the TGN-proximal lysosome is storage of copper for subsequent utilization by the cells in times of copper starvation.

We found that in cells of hepatic origin, ATP7B localizes at endolysosomes besides TGN in basal (physiological) as well as in depleted copper conditions. Interestingly, in hepatic cells but not in other cells of non-hepatic origin. Besides, knocking down ATP7B in HepG2 led to an adverse effect on lysosomal functioning ^67^. In chordates, liver is the main site of nutrient recycling. Copper being a micronutrient is possibly reutilized in the cytosol and other organelles via lysosomal recycling. We hypothesize that in basal physiological conditions, endolysosome-resident ATP7B imports copper in the endolysosomal lumen upon being laden with copper by cytosol localized metallochaperone ATOX1. The less studied copper transporter of the CTR family, CTR2, might be a partner in endolysosomal copper recycling, as it has been shown to export endosomal luminal copper to the cytoplasm. ATP7B and CTR2 works in unison maintain the copper levels in the endolysosome-cytoplasm system. It has been recently shown that the S29L mutation in CTR2 has a relative higher frequency in a cohort of WD patients from China as compared to control individuals ^78^. Interestingly, S29L leads to increased intracellular Cu concentration and Cu-induced apoptosis in cultured cell lines. It would be important to further characterize the functional interplay of ATP7B and CTR2 to understand the underlying reason to localization of ATP7B at the endolysosomal compartments. A comprehensive illustration of recycling of ATP7B that summarizes our study is illustrated in Fig.8D.

## Materials and methods

### Plasmids and antibodies

mEGFP-ATP7B and mKO-ATP7A constructs was available in lab. mKO-ATP7B construct was prepared by cloning ATP7B in mKO-ATP7A construct using MluI and NotI restriction site. The GFP-Arl8b and GFP-RUFY3 constructs were a kind gift from Dr. Amit Tuli, CSIR – Institute of Microbial Technology, India. Ratiometric pHluorin tagged TGN38 C-terminal and dsRed-GalT4 were a kind gift from Prof. Carolyn Machamer, Johns Hopkins University. Mutations on mEGFP-ATP7B were prepared following Q5^®^ Site-Directed Mutagenesis Kit (NEB #E0554) protocol. Preparation of H1069Q mutant is described in Das et al ^53^. Primers used for S653Y mutant are forward-5’TGGAAGAAGTATTTCCTGTGC3’ and reverse-5’CTGCTTTATTTCCATCTTG3’. Plasmid isolation was done using Macherey Nuggel plasmid isolation kit (#MN740490).

### Cell lines and cell culture

HepG2 cells were grown and maintained in complete medium containing low glucose Minimum Essential Medium (MEM) (Himedia #AL047S) supplemented with 10% Fetal Bovine Serum (Gibco #10270-106), 1X Penicillin-Streptomycin (Gibco #15140-122). Similarly, HEK293Tcells were grown and maintained in Dulbecco’s modified Eagle’s medium (DMEM) (Sigma #D6429) supplemented with 10% Fetal Bovine Serum, 1X Penicillin-Streptomycin. For transfection, JetPRIME transfection reagent (Polyplus #114-07) was used as per manufacturer’s protocol.

For copper treatment, 100µM CuCl_2_ was used for 2h, else mentioned otherwise. For creating copper deprived condition, cells were treated with 100 µM BCS or 25 µM TTM for 2 h. For prolong copper chelation, cells were treated with 100 µM BCS for 12 h. For chronic copper chelation, cells were grown in 10 µM BCS for 72 h followed by 100 µM BCS for 2 h. For small molecule treatments, Brefeldin-A (Sigma #B5936), Rapamycin (Sigma #R8781), Cycloheximide (Merck#239763), Bafilomycin A_1_ (Sigma #SML1661) was used 30min prior to any copper treatment, if not mentioned otherwise.

### Immunofluorescence and microscopy

For fixed cell imaging, cells were fixed in 2% PFA in PBS solution for 20 min followed by PFA quenching in 50mM NH_4_Cl solution for 20 min. Blocking was done in 3% BSA (bovine serum albumin) in PBSS (0.075% saponin in PBS). Cells were incubated in primary antibody in room temperature for 1.5 h and in secondary antibody for 30 min. After subsequent washing with PBSS, Coverslips were fixed on glass slides using SIGMA Fluoroshield™ with DAPI mountant. (Sigma #F6057) or ProLong^®^ Gold Antifade Reagent (CST #9071). The solvent for antibody dilution was 1% BSA in PBS. For live cell imaging, cells were seeded on confocal dishes. During imaging cells were kept in imaging media (Phenol red free MEM, Gibco # 51200038; supplemented with 10% FBS, 20mM HEPES and 1% Trolox, Sigma #238813). All images were acquired with Leica SP8 confocal platform using oil immersion 63X objective (NA 1.4) and deconvoluted using Leica Lightning software.

Details of Antibodies used are as follows: ATP7B(Abcam #ab124973, 1:400), TGN46 (Bio-Rad #AHP500G, 1:400), LAMP1 (DSHB #H4A3, 1:200), LAMP2 (DSHB #H4B4, 1:200; Affinity Biosciences #DF4806, 1:400), Golgin-97 (CST #13192, 1:400), Rab7a (SCBT#sc376362, 1:50), GCC2 (Atlas antibodies #HPA035849, 1:400), GM130 (BD Biosciences #610822, 1:400), TGN38 (Novus #NBP103-495, 1:100), EEA1 (BD Biosciences #610457, 1:400), Calnexin (Novus #NB300-518, 1:200; Abcam #ab22595, 1:400), Goat anti-Rabbit Alexa488 (Thermo #A-11034, 1:2000), Goat anti-Rabbit Alexa568 (Thermo #A-11011, 1:2000), Donkey anti-Mouse Alexa 555 (Thermo #A-32773, 1:2000), Donkey Anti-Mouse Alexa405 (Abcam #ab175658, 1:1000), Donkey anti-Sheep Alexa 647 (Thermo #A-21448, 1:2000).

### Immunoblotting

After respective treatment, cells were pelleted down and pellete was dissolved in 100µL of RIPA lysis buffer (10mM Tris-Cl pH 8.0, 1mM EDTA, 0.5mM EGTA, 1.0% Triton X-100, 0.1% sodium deoxycholate, 0.1% SDS, 140mM NaCl, 1X protease inhibitor cocktail, 1mM phenylmethylsulfonyl fluoride (PMSF)) and kept for 30mins on ice. The solution is then sonicated with a probe sonicator (6 pulses, 5sec, and 100mA). Bradford was used for protein quantitation and 20 µg protein was loaded in each well. Protein sample was prepared by adding 4X NuPAGE loading buffer (Invitrogen#NP0007) to a final concentration of 1X and ran on 8% SDS PAGE. This was further followed by semi-dry transfer of proteins onto nitrocellulose membrane (Milipore #IPVH00010). After transfer, the membrane was blocked with 3% BSA in TBST (TBS with 0.1% Tween20) for 3h at RT with mild shaking. Primary antibody incubation was done overnight at 4°C and then washed with 1X TBST (0.01% Tween-20). HRP conjugated respective secondary incubation was done for 1.5 h at RT, further washed and signal was developed by Clarity Max Western ECL Substrate (BioRad #1705062) in ChemiDoc (BioRad).

Details of Antibodies are as follows: ATP7B (1:1500), α-tubulin (Affinity Biosciences#AF7010, 1:20000), LAMP1 (DSHB, 1:400), LAMP2 (1:400), Rab7 (1:500), Atox1 (Merck #WH0000475M1, 1:500), ATP7A (Abcam #ab131400, 1:1000), CTR1 (Abcam #ab129067, 1:4000), Anti-rabbit HRP (CST #7074, 1:6000), Anti-mouse HRP (CST #7076, 1:6000).

### Quantitative Real-time PCR

Cells were scraped from 60mm dishes at 70% confluency using 1 ml of TRIzol™ Reagent (Thermo #15596018). After 5min incubation, 0.2ml of chloroform (SRL #96764) was added and mixed by mild shaking. After that, the mixture was centrifuged at 12,000g for 15min at 4°C. 200µL of the aqueous phase was collected and 500 µL isopropanol (SRL #38445) was added to it and incubated for 10min followed by centrifugation at 12,000g for 10min at 4°C. RNA pellet was washed with 500 µL of 75% ethanol twice followed by centrifugation at 12,000g for 5min at 4°C and then was dissolved in 50 µL DEPC treated water. cDNA was prepared from 1 µg of RNA using Biobharti Super RT MulV kit (Biobharti #BB-E0045). Real-time PCR was performed with SYBR green fluorophore (BioRad #) using BioRad CFX-96 Real Time System. Relative transcript level of all the genes were normalized against HepG2 cells and RLP13A gene was taken as endogenous control. The experiment was performed as per minimum information required for publication of quantitative RT PCR.

Details of primers are as follows: RPL13A_For: GTATGCTGCCCCACAAAACC; RPL13A_Rev: TTCAGACGCACGACCTTGAG; ATOX1_For: GCACGAGTTCTCCGTGGAC; ATOX1_Rev: GGGCCAAGGTAGGAAACAGC; ATP7A_For: GAAGAGGTCGGACTGCTGTC; ATP7A_Rev: CCTTAGTAATGCCAACCTGAGAAGC; CTR1_For: TCTCACCATCACCCAACCAC; CTR1_Rev: AAAAGCTCCAGCCATTTCTCC. All primers are from 5’ to 3’ direction.

### Sample preparation for ICP-OES and ICP-MS

At 70% confluency, HepG2 cells, seeded on 60mm dishes, were treated with respective treatment conditions. After that, cells were quickly washed with 1X DPBS (Gibco #14200075) for 10 times and then were scrapped using cell scrapper. Cells were collected by centrifuging at 2000 rpm for 5 min. For cellular copper measurement of HepG2 in different copper condition, the cell pellets were dissolved in 100µl of 1X DPBS. After that, 2.5 million cells were dissolved into 100 µl of ICP-OES grade 65% HNO_3_ acid (Merck #1.00441.1000) and was kept at 95°C for overnight. For cellular copper measurement of different cells and prolong copper chelation, cell pellet was dissolved in 100 µl of 1XDPBS followed by sonication with a probe sonicator (6 pulses, 5sec, and 100mA). Bradford was used for protein quantitation and 500 µg protein were digested overnight. Next day, 5ml of deionized water was added to it and were filtered through a 0.45-micron filter. Copper calibration is done by acid digestion of copper foil (procured from Alfa Aeasar) in 10 mL suprapure HNO_3_ for 1 h. (MWD conditions: Power=400 W; Temperature=100^0^C; Hold time= 1 h). From the obtained solution, different solutions of varying copper strengths were used for calibration. Copper concentration was determined using an iCAP 6500 ICP-OES (Thermo Scientific) or a Xseries 2 ICP-MS (Thermo Scientific).

### Cellular fractionation using sucrose gradient

For cellular fractionation, the protocol were modified from Carosi et al ^79^. Cells were seeded in 100mm dishes. At 70% confluency, cells were washed in 1X DPBS and scrapped in 250µL of homogenization buffer (250mM sucrose, 1mM EDTA, 1X protease inhibitor cocktail, 1mM PMSF) and lysed by passing through 30-gauge needle for 100 times. After that cells were centrifuged at 1000g for 10min at 4°C. The lysate was collected and adjusted to 1ml of 25% sucrose and 1mM EDTA solution. In a decending order of concentration the following gradient of sucrose solutions were loaded: 2.5ml of 45% sucrose solution followed by, 5.3ml of 35% and 4.2ml of 25% sucrose solution in a. 1ml of cell lysate was added at top and it was centrifuged at 2,00,000g for 2 h at 4°C. 1ml of fractions were collected and immunoblotting was done accordingly.

Details for antibodies are as follows: ATP7B (1:1500), LAMP2 (1:400), GGA2 (BD Transduction Laboratory #612612, 1:2000), TGN46 (1:1000), EEA1 (1:1000), Rab7 (1:500), Anti-rabbit HRP (1:6000), Anti-mouse HRP (1:6000). For TGN46, Rabbit Anti-Sheep IgG(H+L)-AP (Southern biotech #6150.04, 1:2000) was used along with NBT (#S380C) and BCIP (#S381C) (All were kind gift from Dr. Bidisha Sinha, IISER Kolkata).

### Time-lapse fluorescence microscopy

HepG2 cells were seeded on confocal dishes (SPL) and were co-transfected using jetPRIME (Polyplus) transfecting reagent as per the manufacturer’s protocol. For marking acidic compartment, cells were incubated with 0.5 mg/ml lysine-fixable dextran–Alexa Fluor 647 (Thermo #D22914) for 4 h followed by chasing in conjugate-free medium for 20 h as previously described ^80^. Cells were kept in imaging media (phenol red-free MEM supplemented with 2%FBS, 20mM HEPES and 1mM TROLOX) during imaging. Images were acquired using Leica SP8 confocal platform using oil immersion 63X objective (NA 1.4) and deconvoluted using Leica Lightning software. Maximum Intensity Projection (MIP) was made in LAS-X software provided by Leica and videos were processed using Cyberlink Powerdirector.

### Proximity ligation assay

HepG2 cell, seeded on glass coverslips, was fixed with 4% PFA after respective treatments and permeabilized with 0.1% saponin. After that, the proximity ligation assay was performed following the manufacturer’s protocol (Duolink™ PLA Technology, Sigma). Cells was mounted in SIGMA Fluoroshield™ with DAPI mountant. (Sigma#F6057). Images was acquired with Leica SP8 confocal platform using oil immersion 63X objective (NA 1.4) and deconvoluted using Leica Lightning software. Details for antibodies are as follows: LAMP2 (1:100), GCC2 (1:200), ATP7B (1:200), Vps35 (SCBT#sc-374372, 1:75).

### Flow cytometry Assay

At 70% confluency, HepG2 cells, seeded on 12 well plates were treated with respective treatments. After treatment, cells were washed in 1X DPBS. Lysosome was stained with 100nM LysoTracker red DND 99 (Thermo #L7528) and cells were incubated for 30 min. LysoTracker was a kind gift from Dr. Sayam Sen Gupta (IISER Kolkata). Cells were washed with 1X DPBS and kept in 50 µL Trypsin for 10 min. After that, cells were suspended in 350 µL of FACS buffer (1XDPBS, 2% FBS, 25mM HEPES, 2mM EDTA) and was analysed by flow cytometry BD LSRFortessa (BD Biosciences). 10,000 cells were analysed for each condition. FCS Express software (version7.14.0020) was used for data analysis. Mean intensity value were used for quantification.

### Lysosomal exocytosis Assay (Hexosaminidase assay)

HepG2 cells were seeded in a 12-well plate. At 60% confluency, cells were washed with 1XDPBS and kept in 200μl of phenol red free media (Gibco #51200038, along with 10% FBS) for the treatment duration. Following incubation, media was collected and centrifuged at 12,000g for 15min at 4°C and the supernatant was collected. Remaining cells were lysed in 500μl lysis buffer (HEPES 25mM, NaCl 150mM, 0.5% Triton-100, 1X protease inhibitor cocktail) using probe sonicator (6 pulses, 5sec, and 100mA) followed by centrifugation 12,000g for 15min at 4°C and the supernatant was collected. For Hexosaminidase assay, 75 µL of media supernatant (denoted as SN) and cell lysate (denoted as L, 10 µL of cell lysate and 65 µL of 1X PBS) was added in 96-well plate in triplicate. Media and Lysis buffer control was taken as blank (denoted as bl). 75μl of 3mM 4-nitrophenyl N-acetyl-β-d-glucosaminide (Sigma #N9376) for in 0.1M citrate buffer (0.1M sodium citrate, 0.1M citric acid, pH 4.5) 30 min at 37°C. Reactions were stopped by adding 50μl of borate buffer (100mM boric acid, 75mM NaCl, 25mM sodium borate, pH 9.8) and the absorbance was measured at 405nm. Relative enzyme activity was calculated from the E_SN_/E_L_ Ratio after substracting the blank E_bl_ and then normalized to Basal.

### Immunofluorescence of mouse liver

Mouse liver embedded in Tissue freezing media was sectioned using Cryotome. Tissue sections were deparaffinized at 60-65°C keeping on hot plate then and rehydrated in xylene, 100% ethyl alcohol, 70% ethyl alcohol, distilled water, and 1X PBS solution. Antigen unmasking was performed at 95°C in antigen unmasking solution for 15 min. Sections outlined by hydrophobic PAP pen and incubated in blocking serum for 1 h at 37°C in a humidified chamber. After that, tissue sections were incubated in primary antibody overnight at 4°C. Secondary antibody was carried on for 2hr. Sections were mounted in SIGMA Fluoroshield™ with DAPI mountant (Sigma#F6057). Images was acquired with Leica SP8 confocal platform using oil immersion 63X objective (NA 1.4) and deconvoluted using Leica Lightning software. Antibody for ATP7B was raised in rabbit against amino-terminal of ATP7B from Bio Bharati Life Science Pvt. Ltd. Details of antibody used are as followed: anti-ATP7B amino terminal (1:50), LAMP2 (DSHB, 1:40), Goat anti-Rabbit Alexa 586 (Thermo, 1:800), Donkey anti-Mouse Alexa 488 (Thermo, 1:800).

### Immuno-EM of endogenous ATP7B in HepG2

The samples for electron microscopy (EM) were prepared as described earlier (Polishchuk et al 2003; PMID: 12937271). Briefly, the cells were fixed in fixative containing 4% paraformaldehyde and 0.05% glutaraldehyde for 10 min at RT. The cells were then permeabilized and stained with ATP7B antibody ((rabbit polyclonal, Abcam, Cambridge, UK, ab124973) followed by second antibody labelled with nanogold. The nanogold signal was then amplified with gold enhancement (Nanoprobes). The samples were then dehydrated, embedded in epon and sectioned (60-70nm thin sections). The sections were then imaged using FEI Tecnai 12 120kV microscope.

### Image analysis and statistics

Images were analyzed in batches using ImageJ ^81^, image analysis software. For colocalization study, Colocalization_Finder plugin was used. ROIs were drawn manually on best z-stack for each cell. Mender’s colocalization coefficient (MCC) ^82^ and Pearson correlation coefficient(PCC) ^83,84^ were used for quantifying colocalization. Merged image of LAMP2 channel and TGN46 channel was obtained in greyscale which was used to obtain the total lysosomal ATP7B. For counting PLA positive spots, images were converted to binary using default threshold and Analyze Particles tool was used with a size cutoff of >5 pixel along with circularity <0.9 to avoid false positive counting due to non-specific staining. For the box plots, the box represents the 25th to 75th percentiles, and the median in the middle. The whiskers show the data points within the range of 1.5 × interquartile range (IQR) from the first and third quartile. Non-parametric tests for unpaired datasets (nonparametric Mann–Whitney U test/Wilcoxon rank-sum test) were performed for all the samples; ∗*p* < 0.05, ∗∗*p* < 0.01, ∗∗∗∗*p* < 0.0001; ns, not significant. For statistical analysis and plotting, ggplot2 and ggpubr package was used in R v-4.2.1 ^87^. ImageJ macro codes for image analysis are available at https://github.com/saps018/tgn-lcs

## Acknowledgments

This work is supported by DBT-Wellcome Trust India Alliance Fellowship (IA/I/16/1/502369), and Core Research Grant (CRG/2021/002150) from SERB, Department of Science and Technology (DST), Government of India, and IISER-K intramural funding to AG. SM is supported by a pre-doctoral fellowship from Council of Scientific and Industrial Research (CSIR), India. RP is supported by a pre-doctoral fellowship from Department of Biotechnology, Govt. of India. The Pre-doctoral fellowship for Ruturaj is supported by Intramural Institute funding (IISER-K). We thank Amit Tuli (IMTECH, Chandigarh), Enrique Rodriguez-Boulan, Ryan Schreiner (Weill Cornell Medical College) and Carolyn Machamer (Johns Hopkins University) for sharing reagents with us. We would also like to thank Dr. Santosh Ch Das (Department of Earth Sciences, IISER Kolkata), the ICP-MS and ICP-OES facility, IISER-Kolkata and imaging facility, ILS Bhubaneswar for helping us with our experiments. We thank the Euro-BioImaging facility at the IEOS (CNR), Naples for help with microscopy experiments. The facility is supported by grants from MIUR, the Government of Italy and also the grants PON-IMPARA, POR-CIRO and SEELIFE. We thank Dr. Fareeha Saadi (Washington University School of Medicine) for helping with mouse liver immunofluorescence.

## Author contributions

AG and SM designed the experiments and wrote the manuscript. SM, MP, RP, Ruturaj and SD did the experiments and analysed the data. SM wrote the codes and analysed the data. Ruturaj helped with time-lapse imaging. TG conducted the Flow cytometry experiments and analysed the data. MP performed the immune-EM and analysed the data. All authors reviewed the results and approved the final version of the manuscript.

**The authors declare no competing financial interests.**

**The manuscript does not require ethics committee approval at the institution.**

## Data availability statement

The authors confirm that the data supporting the findings of this study are available within the article and/or its supplementary materials.

## Supplementary figure legends

**Figure S1:** Immunofluorescence image of ATP7B (green) in HepG2 cell line co-stained with endolysosomal marker LAMP2 (red) and TGN marker TGN46 (blue) in **(A) (i)** low copper (20µM Cu) and **(ii)** high copper (250µM Cu) treatments as well as **(B)** copper chelated (25 µM TTM for 2h). The marked area on ‘merge’ panel is enlarged in ‘inset’. The arrowheads show colocalization between ATP7B-LAMP2, the asterisk (*) shows colocalization between ATP7B-TGN46. **(C)** Immunofluorescence image of ATP7B (green) in HepG2 cell line co-stained with endolysosomal marker LAMP2 (red) and TGN marker TGN46 (blue) shows presence of ATP7B on both the compartments, irrespective of cellular copper condition i.e. basal, high copper (20µM Cu, 100µM Cu and 250µM Cu) as well as copper chelated conditions (25µM TTM, 100µM BCS). The marked area on ‘merge’ panel is enlarged in ‘inset’. The white arrow shows colocalization between ATP7B-LAMP2, the asterisk shows colocalization between ATP7B-TGN46. Scale bar: 5µm. **(D)** The fraction of ATP7B in TGN46 and LAMP2 in different copper condition are demonstrated by box plot with jitter points. The black * shows the comparison of colocalization between ATP7B-TGN46 and ATP7B-LAMP2. The green * shows the comparison of ATP7B-TGN46 colocalization under different condition with the basal. The orange * shows the comparison of ATP7B-LAMP2 colocalization under different condition with the basal. Sample size (N) for TTM: 46, BCS: 33, Basal: 36, Cu20: 23, Cu100: 33, Cu250: 67. **(E)** Immunofluorescence image of ATP7B (green) in HepG2 cell line co-stained with endolysosomal marker LAMP2 (red) and TGN marker TGN46 (blue) shows presence of ATP7B upon 12h copper chelation using 100 µM BCS. The marked area on ‘merge’ panel is enlarged in ‘inset’. The white arrow shows colocalization between ATP7B-LAMP2, the asterisk shows colocalization between ATP7B-TGN46. Scale bar: 5µm. **(F)** The PCC of ATP7B-LAMP2 in different copper condition are demonstrated by box plot with jitter points. Sample size (N) for Basal: 66, 12h BCS: 55, Chronic BCS: 87. **(G)** Immunoblot of ATP7B after different copper treatment. α-tubulin has been used as a housekeeping control. The fold changes of ATP7B abundance normalized against α-tubulin has been mentions. **(H)** Immunofluorescence image of ATP7B (green) in HepG2 cell line co-stained with EEA1 or Rab7 (red) and TGN marker TGN46 (blue) shows absence of ATP7B on both the compartments (EEA1 or Rab7) **(I)** The PCC of ATP7B with EEA1 or Rab7 in basal condition are demonstrated by box plot with jitter points. [Scale bar: 5µm.]

**Figure S2:** Immunofluorescence image of **(A)** endogenous ATP7B (green) in HEK293T cell line and **(B)** ectopically expressed mEGFP-ATP7B (green) in HeLa cells co-stained with lysosomal marker LAMP2 (red) and TGN marker TGN46/GCC2 (blue) shows different localization pattern of ATP7B that hepatocytes. The marked area on ‘merge’ panel is enlarged in ‘inset’. The arrowhead shows colocalization between ATP7B-LAMP2, the asterisk shows colocalization between ATP7B-TGN46. **(C)** Ectopically expressed mKO-ATP7A (red) in HepG2 cell co-stained with ATP7B (green) and TGN46 (blue) under Cu treated condition. The arrowhead marks the membrane localization of ATP7A under copper treated condition. **(D)** Comparison of copper accumulation in HepG2, HUH7, HEK293T and HeLa cells (n = 9) in basal condition shows higher copper accumulation in HepG2 cell w.r.t. other cells. For side-by-side comparison, prolong copper chelated conditions (12h and chronic chelation, 72h using BCS) shows comparable copper accumulation. Copper concentrations were measured in parts per billion (ppb, mean ± SD). **(E)** RT-PCR of ATOX1, ATP7A and CTR1 in HepG2, HUH7, HEK293T, and HeLa is represented as bar plot (mean ± SEM). ATOX1 and CTR1 levels are comparable in all the cells. ATP7A level in much higher in HEK293T than other cells. **(F)** Immunoblot of ATOX1 (∼8kDa monomer, ∼16kDa dimer), ATP7A (∼170kDa) and CTR1 (∼25kDa) in HepG2, Huh7, HEK293T, HeLa shows similar trend as seen in RT-PCR data. Loading control: α-tubulin [Scale bar: 5µm.]

**Fig S3:** Immunofluorescence of ATP7B (green) co-stained with different TGN-endolysosome combinations i.e. **(A)** LAMP1(red)-TGN46(blue), **(B)** LAMP1(red)-Golgin97(blue), **(C)** LAMP2(red)-TGN38(blue) shows presence of ATP7B in TGN marker –endolysosome marker colocalizing regions (marked with arrowhead) in HepG2. **(D)** Though, ATP7B was absent in Rab7(red)-TGN46(blue) colocalization region (marked with arrow). **(E)** Presence of ATP7B in TGN-proximal lysosome in Huh7 cell is marked by co-staining ATP7B (green), LAMP2 (red) and TGN46 (blue). Immunofluorescence of ATP7B (green) co-stained with **(F)** Calnexin(red)-TGN46(blue), **(G)** LAMP2(red)-Calnexin(blue) shows absence of ATP7B in TGN-ER and lysosome-ER colocalizing region. Intensity profile upon the drawn line is represented on the extreme-right panel of each figure. [Scale bar: 5µm.]

**Fig S4: (A)** GLCS is marked with LAMP1 (green) and GM130 (red) colocalization, (marked with arrowhead). TGN-lysosome proximity region is marked with LAMP1 (green) and TGN46 (blue) colocalization, (marked with asterisk). **(B)** ATP7B (green) co-stained with LAMP1 (red) and cis-Golgi marker GM130 (blue) shows absence of ATP7B in the GLCS (marked by colocalizing regions of LAMP1 and GM130). **(C)** Colocalization between TGN46-LAMP2 shows increased colocalization in BFA treated cells than control (ct). Sample size (N) are ct: 43, BFA: 63. **(D)** Flow cytometry analysis shows decrease in Lysotracker intensity upon mild Bafilomycin A1 treatment (40nM for 30min). Data is normalized against ct and demonstrated as bar-plot (mean ± SD). **(E)** Immunofluorescence of ATP7B (green) co-stained with TGN46 (blue) and LAMP2 (red) shows no change trafficking of ATP7B in mild Bafilomycin-A_1_ pre-treatment (40nM for 30min), both in basal (top) and Cu (bottom) treated conditions (marked with arrowhead). Marked area in the first panel is enlarged in the next panels. **(F)** The PCC between LAMP2-TGN46, ATP7B-TGN46 and ATP7B-LAMP2 in basal and copper treated conditions in control, ct (green) and Bafilomycin-A1 (orange) pre-treated conditions are demonstrated by box plot with jitter points. Sample size (N) for ct are Basal: 57, Cu: 51; for Baf-A treated are Basal: 40, Cu 49. [Scale bar: 5µm.]

**Fig S5: (A)** Cells ectopically expressing GFP-Arl8b and GFP-RUFY3 are shown in magenta. **(B)** Proximity ligation assay shows increase PLA positive punctate (red) in Rapamycin treated conditions **(C)** Quantitation of PLA positive punctate in basal copper and Rapamycin treated condition is demonstrated by box plot with jitter points. Sample size (N) are Basal: 42 Cu: 20, Rapamycin: 37. **(D)** Immunostaining of ATP7B (green) co-stained with LAMP2 (red) and TGN46 (blue) in HepG2 in control and CHX treated conditions in Cu250>BCS 30min treatment. **(E)** The fraction of ATP7B present in LAMP2 and TGN46 in control (ct) and CHX treated cells under different copper conditions are demonstrated by box plot with jitter points. **(F)** The PCC between ATP7B-LAMP2 and ATP7B-TGN46 in ct and CHX treated cells under different copper conditions are demonstrated by box plot with jitter points. Sample size (N) for (E) and (F) in basal condition are basal: 94, Cu250: 108, Cu_BCS30m: 104, Cu_BCS2hr: 99. Sample size for CHX treated condition are basal: 57, Cu250: 84, Cu_BCS30m: 100, Cu_BCS2hr: 103. [Scale bar: 5µm.]

