## Supplementary figures and images for "Copper-independent lysosomal localization of the Wilson disease protein ATP7B"

### Supplemental

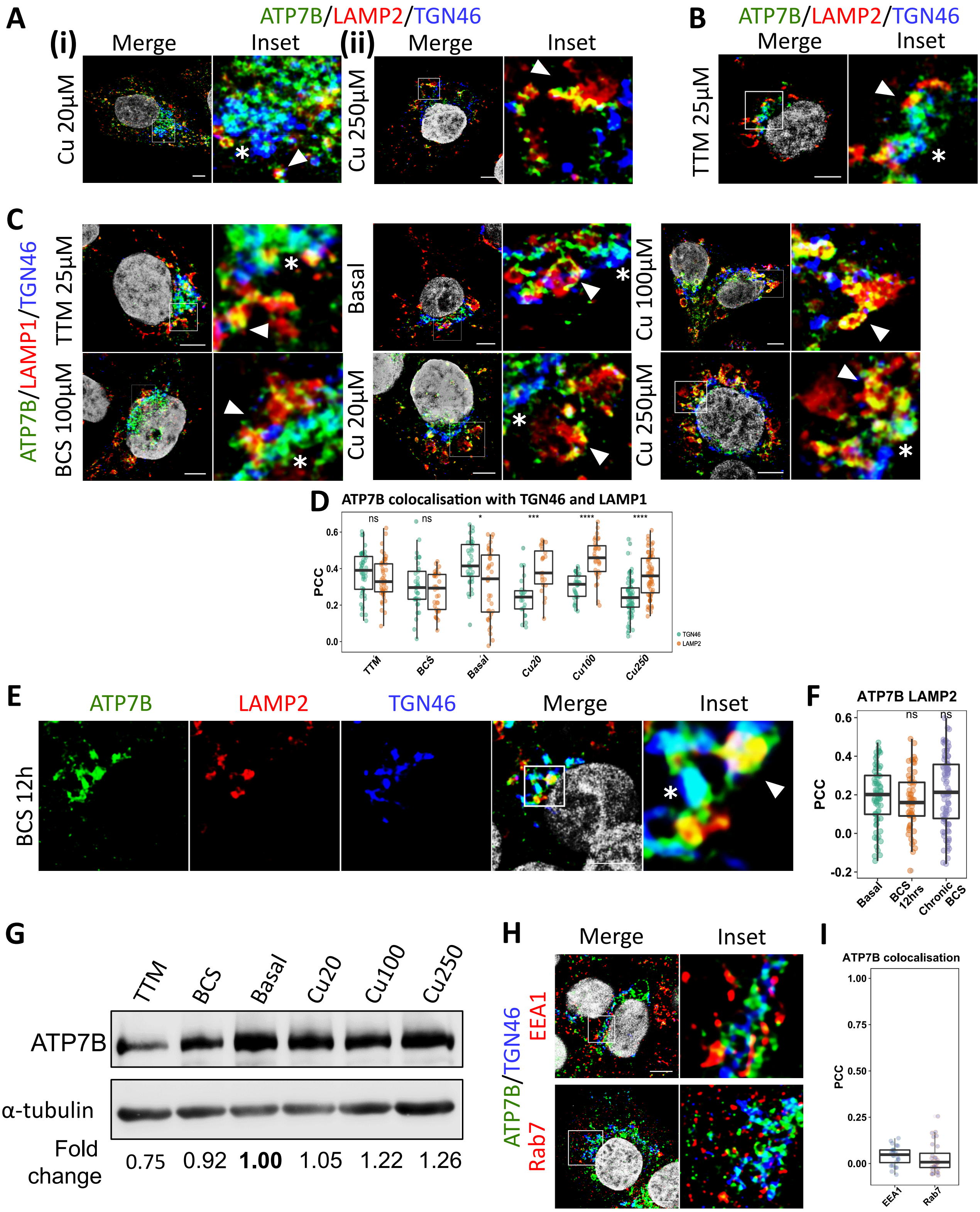

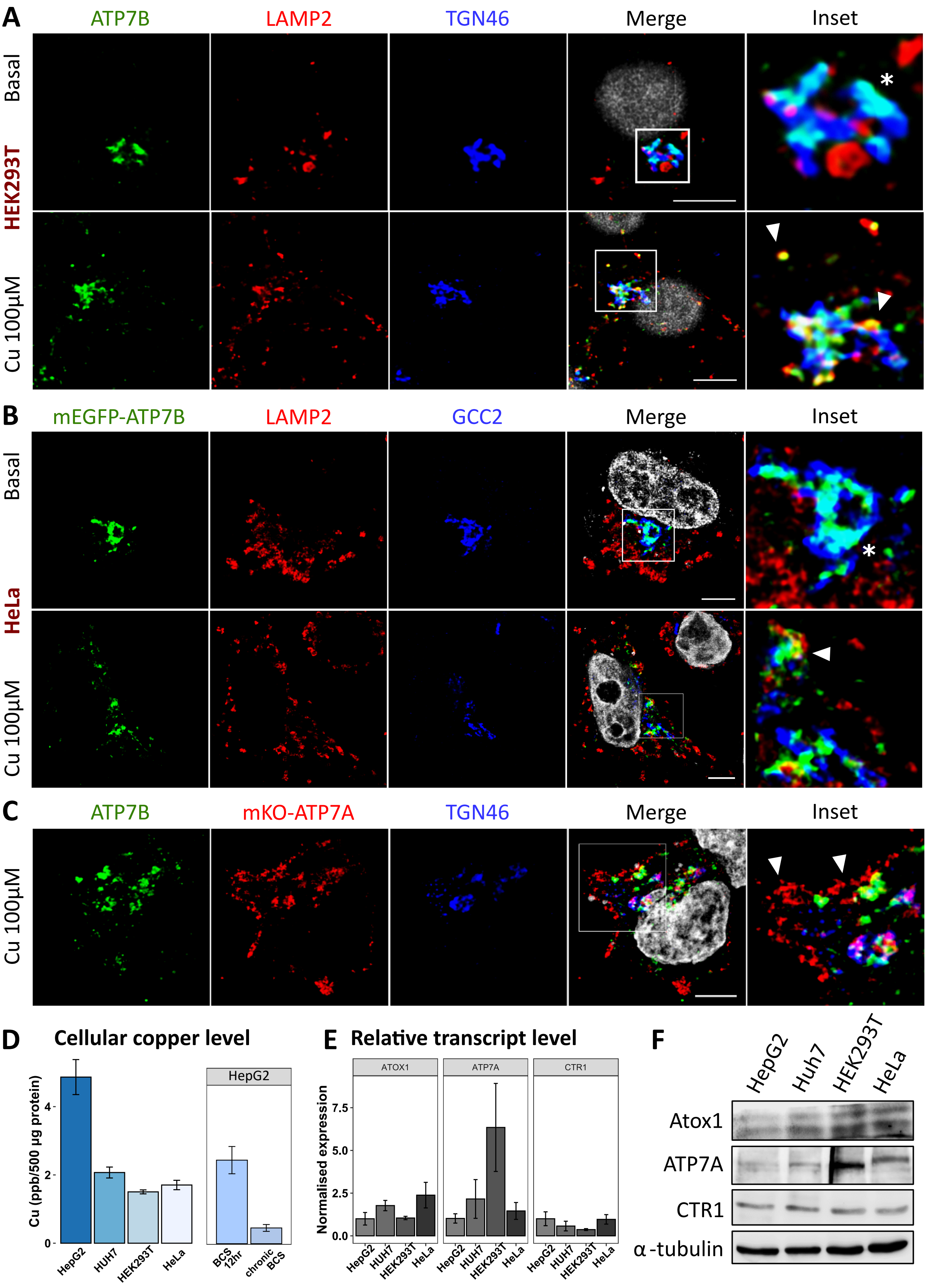

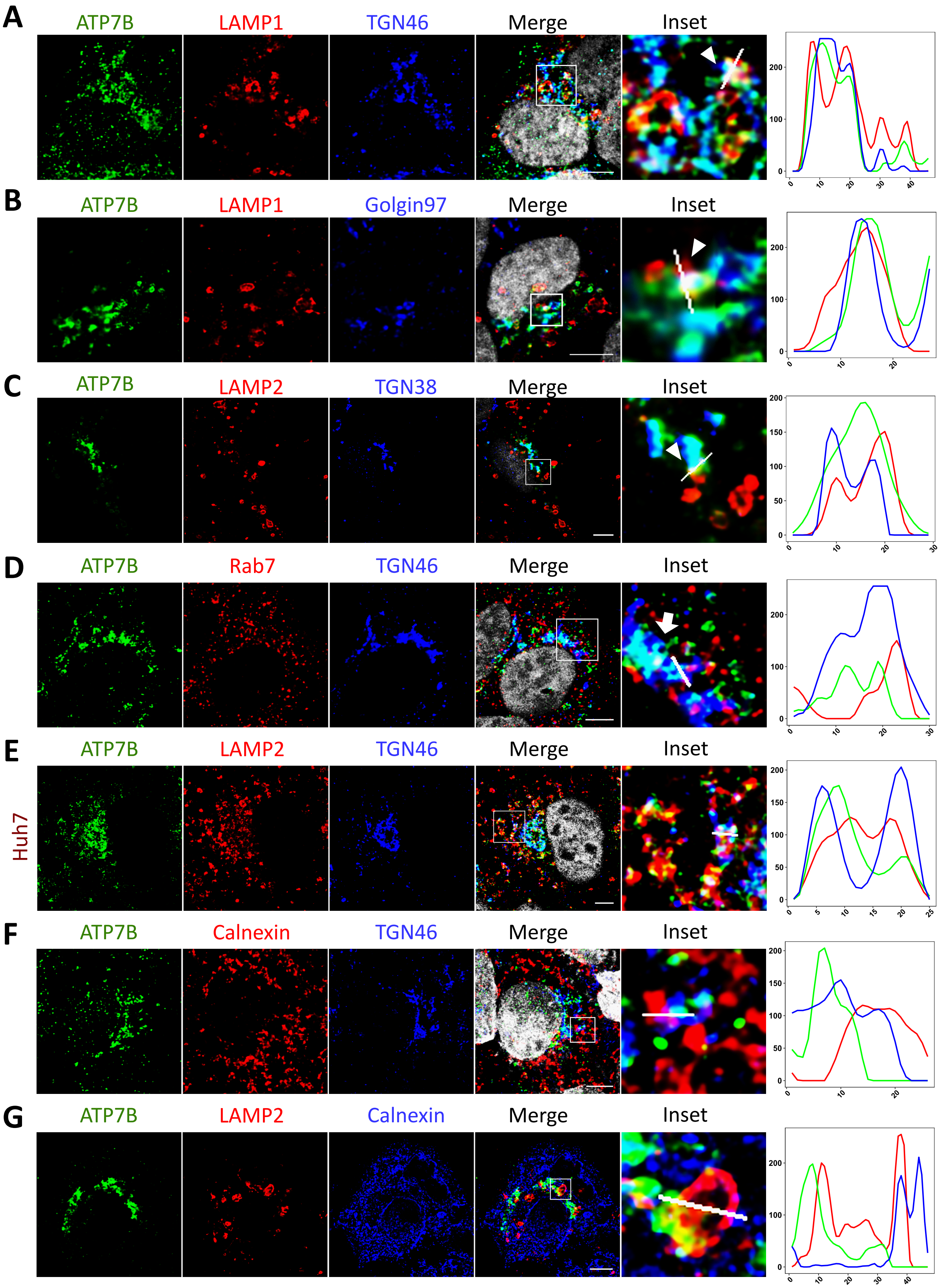

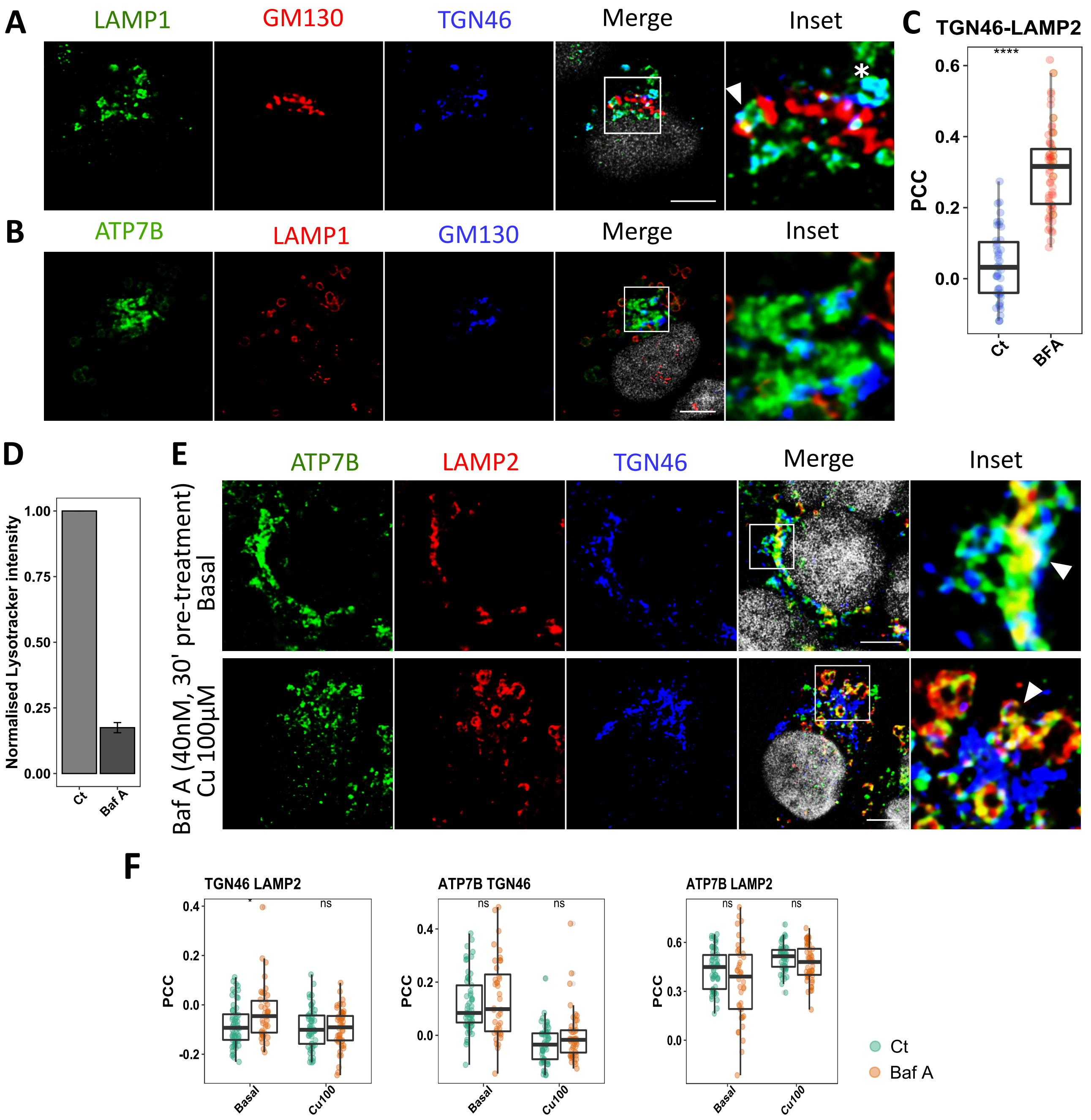

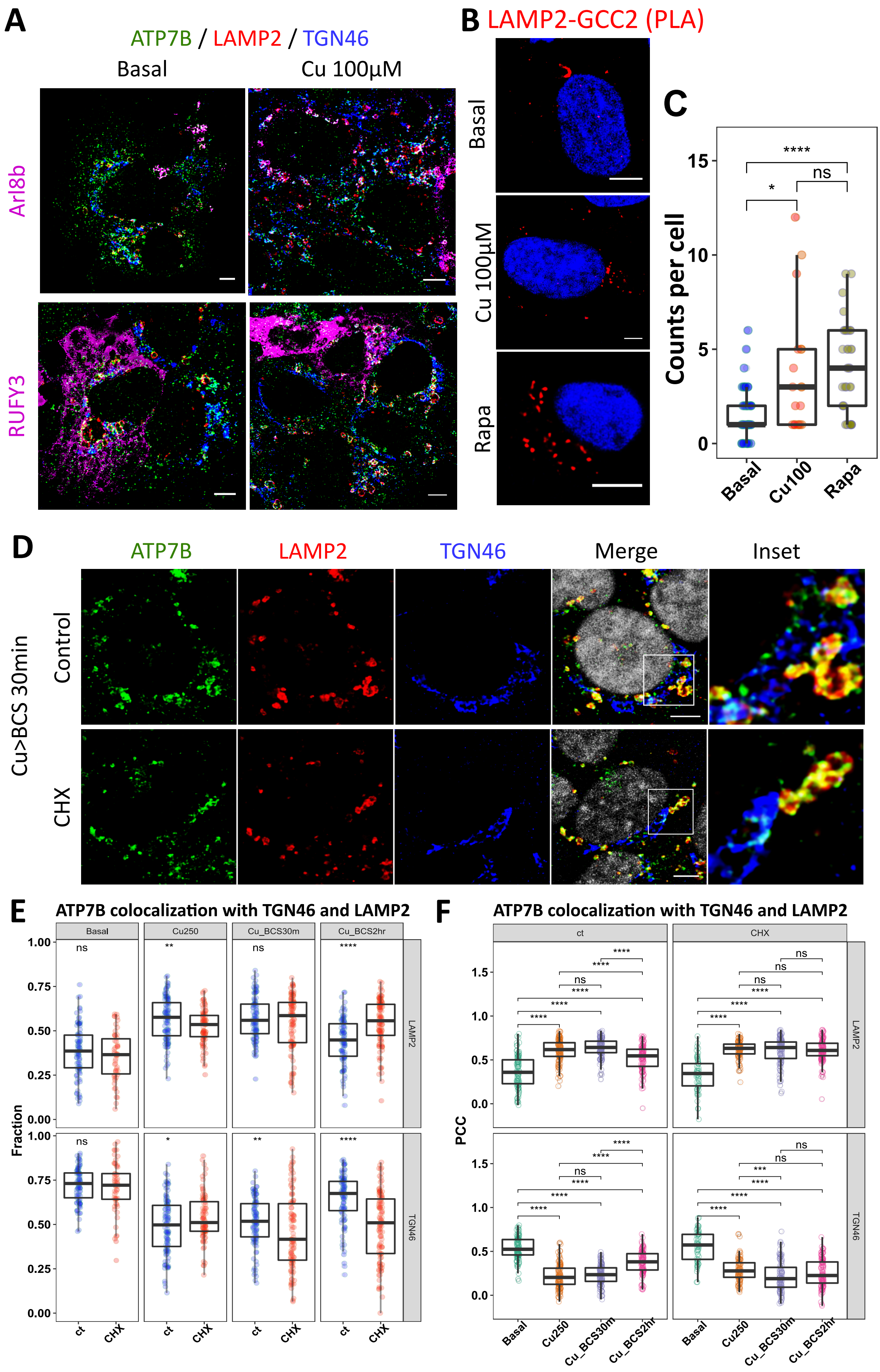
